# Diagnostic tool to identify and treat DNA repair deficient gastroesophageal adenocarcinomas

**DOI:** 10.1101/2022.07.14.500118

**Authors:** Aurel Prosz, Pranshu Sahgal, Clare X. Morris, Zsofia Sztupinszki, Judit Börcsök, Miklos Diossy, Viktoria Tisza, Sandor Spisak, Orsolya Rusz, Istvan Csabai, Brandon M. Huffman, Harshabad Singh, Jean-Bernard Lazaro, Michael Cecchini, James M. Cleary, Zoltan Szallasi, Nilay S. Sethi

## Abstract

**Background and aims:** DNA repair deficiency is a common feature of cancer. Homologous recombination (HR) and nucleotide excision repair (NER) are the two most frequently disabled DNA repair pathways in solid tumors. HR deficient breast, ovarian, pancreatic and prostate cancers respond well to platinum chemotherapy and PARP inhibitors. However, the frequency of DNA repair pathway deficiency in gastric and esophageal adenocarcinoma (GEA) still lacks diagnostic and functional validation. Furthermore, whether DNA repair deficient GEA have enhanced responsiveness to platinum chemotherapy and sensitivity to PARP inhibitors is not well characterized.

**Methods:** Using whole exome and genome sequencing data, we measured various HR deficiency-associated mutational signatures in patient specimen of gastric, esophageal and colorectal cancer specimens and gastric cancer cell lines. Gold-standard immunofluorescence assays were used to confirm HR and NER deficiency in cancer cell lines. The relationship between PARP inhibitor treatment and tumor response was evaluated in patients with gastric cancer. Drug sensitivity was determined using standard *in vitro* cell culture assays. Single-cell RNA-sequencing was performed to evaluate gastric cancer response to commonly used chemotherapeutics.

**Results:** We found that a significant subset of GEA, but very few colorectal tumors, show evidence of HR deficiency by mutational signature analysis (HRD score). Gastric cancer cell lines with high HRD mutational signature scores demonstrated functional HR deficiency by RAD51 assay and increased sensitivity to platinum and PARP inhibitors. There was a positive association between HRD scores and tumor response in patients with gastric cancer treated with a PARP inhibitor on a clinical trial. A gastric cancer cell line with strong sensitivity to cisplatin showed HR proficiency but exhibited NER deficiency by DDB2 proteo-probe assay. Single-cell RNA-sequencing revealed that, in addition to inducing general apoptosis, cisplatin treatment triggered ferroptosis in a NER-deficient gastric cancer, which may explain the outlier sensitivity.

**Conclusion:** A subset of upper gastrointestinal tumors have genomic features of HR and NER deficiency and therefore may be more likely to benefit from platinum chemotherapy and PARP inhibition.

## INTRODUCTION

Platinum agents are essential components of chemotherapy regimens for the treatment of gastrointestinal cancers. There are several biological mechanisms rendering solid tumors sensitive to platinum-based treatments including DNA repair pathway aberrations, such as homologous recombination (HR) deficiency or nucleotide excision repair (NER) deficiency. Platinum agents are highly mutagenic and generate both intrastrand and interstrand DNA lesions. The HR repair pathway corrects DNA double-strand breaks created by lesions such as platinum-induced interstrand crosslinks. Alterations in HR genes are associated with increased platinum sensitivity in multiple tumor types including breast and ovarian cancer^1, 2^. HR deficiency is likely to be present in GEA since mutations in key HR genes, albeit with low frequency, have been detected, for example, in gastric adenocarcinoma^3, 4^.

In addition to driving sensitivity to platinum-based agents, HR deficiency forms a synthetic lethal relationship with small molecule inhibitors of poly (ADP ribose)-polymerase (PARP), and PARP inhibitors are now FDA approved for HR-deficient breast, ovarian, prostate, and pancreatic tumors. Initial clinical trials of PARP inhibitors in gastric cancer showed mixed results, with a second-line regimen of Olaparib/paclitaxel showing benefit compared to paclitaxel/placebo in a molecularly unselected population, as well as in ATM deficient patients, in a phase 2 but not in the confirmatory phase 3 GOLD trial ^5, 6^. Suboptimal patient selection may have contributed to the disappointing results seen in the phase 3 study. Patients on the GOLD trial had progressed on first-line chemotherapy, which typically includes platinum chemotherapy; whereas a key biomarker for PARP inhibitor sensitivity in ovarian and pancreatic cancer patients has been platinum sensitivity^7^. In addition, it is widely agreed that new genomic markers of PARP inhibitor sensitivity in GEA are needed, and a goal of ongoing clinical trials is to identify and molecularly define the subpopulation of GEA that are sensitive to PARP inhibition (NCT03008278, NCT01123876).

HR deficiency is found in tumors without canonical HR gene mutations, and such cases are detectable by the presence of HR deficiency-associated mutational patterns (signatures). HR deficiency induces multiple types of genetic alterations ranging from single nucleotide variations to large scale genomic rearrangements. The presence and frequency of these events can be used to calculate clinically applicable mutational signatures, such as the HRD score^8^. High levels of these HR deficiency indicators are associated with better response to PARP inhibitor therapy in ovarian cancer^9^.

Here, we applied a validated genomic signature of HR deficiency, analogous to the FDA approved HRD score, to characterize the landscape of HR deficiency in gastrointestinal cancers. We find that a significant fraction of GEA have elevated levels of HR deficiency signatures and that the frequency of HR deficiency in GEA is higher than predicted based on *BRCA1/2* mutational status alone. We unexpectedly found a human GEA cell line model with outlier cisplatin sensitivity exhibited NER deficiency. We then applied our recently published complex mutational signature of NER deficiency^10^ to the next-generation sequencing data and found that NER deficient cases are likely present in GEA. To characterize cisplatin response in GEA NER deficiency, we performed single-cell transcriptional profiling on our human GEA NER deficient cell line model following chemotherapeutic stress, nominating ferroptosis as an additional mechanism of cancer cell death.

## MATERIALS AND METHODS

### Genotyping

Germline and somatic variants in WGS samples were downloaded from the ICGC data portal. Whole exome somatic and germline vcf files were downloaded from the TCGA data portal. The high fidelity of the reported germline and somatic variants was ensured by the application of the tools default filters (FILTER == “PASS”). The high fidelity of the reported germline and somatic variants was ensured by the application of additional hard filters on the somatic samples: TLOD ≥ 6, NLOD ≥ 3, NORMAL.DEPTH ≥ 15, TUMOR.DEPTH ≥ 20, TUMOR.ALT ≥ 5, NORMAL.ALT = 0, TUMOR.AF ≥ 0.05. The pathogenicity of the variants was assessed by Intervar (version 2.0.2) which classifies variants into five categories: “Benign”, “Likely Benign”, “Uncertain significance”, “Likely Pathogenic” and “Pathogenic”. Mutations in exonic regions that were not synonymous SNVs and classified as “Pathogenic” or “Likely Pathogenic” were considered as deleterious.

In order to estimate tumor cellularity and ploidy and to infer allele-specific copy number (ASCN) profiles Sequenza was used. The fitted models were in the ploidy range of [1, 7] and cellularity range of [0, 1]. When the predictions of a fitted model were significantly different from the 2 expected ploidy and cellularity values, an alternative solution was selected manually. If the copy numbers of either the A or B alleles dropped to zero within the coordinates of a gene, then an LOH event was registered. The final genotypes were determined as: Wild type: no pathogenic or likely pathogenic germline or somatic mutation(s). Wild type with LOH: no pathogenic or likely pathogenic germline or somatic mutation(s), but an LOH event occurred. Heterozygote mutant: a pathogenic or likely pathogenic germline or somatic mutation is present, but no LOH. Heterozygote mutant with LOH: a pathogenic or likely pathogenic germline or somatic mutation is present and an LOH event occurred. Homozygote mutant: an identical germline or somatic mutation is present in both alleles. Compound heterozygote mutant: two different germline and/or somatic mutations are present in both alleles.

### Mutational signature extraction

Single base substitution (SBS) signatures were extracted with the help of the deconstructSigs R package which determines the linear combination of pre-defined signatures that most accurately reconstructs the mutational profile of a single tumor sample. The selected signatures, the linear combination of which could lead to the final mutational catalog, were confined to those, that were reported to be present in both gastric, esophageal and colorectal cancer according to the Catalogue of Somatic Mutations in Cancer (COSMIC) (https://cancer.sanger.ac.uk/cosmic/signatures_v2), and signature 3 were also extracted along with them. After evaluation of a sample’s signature composition, its mutational catalog was reconstructed, and the cosine of the angles between the 96-dimensional original and reconstructed vectors was calculated (cosine similarity).

Structural variants (SVs) for germline and somatic samples were downloaded from the ICGC data portal. The resulting structural variants in each sample were mapped to a 32-dimensional rearrangement signature (RS) catalog described in breast cancer (M). The previously identified matrix of rearrangement signatures (P) was downloaded from the following link: https://static-content.springer.com/esm/art\%3A10.1038\%2Fnature17676/MediaObjects/41586_2016_BFnature17676_MOESM47_ESM.zip. As previously, the M and the P matrices were used in a non-negative least-squares problem to estimate the matrix of exposures to mutational processes (E).

For the gastric cell line samples, the variants were obtained from the DepMap portal (https://depmap.org/portal/download/), and the mutational signature and genomic scar score analysis was performed the same way as the patient samples. For the genotyping, the non-conserving mutations were excluded.

### Genomic scar scores

Three independent DNA-based measures of genomic instability using single nucleotide polymorphism (SNP) arrays were developed on the basis of homologous recombination deficiency (HRD)-associated loss of heterozygosity (HRD-LOH), telomeric allelic imbalance (HRD-TAI) and large-scale state transition (HRD-LST). The HRD-LOH score is the number of 15 Mb exceeding LOH regions that do not cover the whole chromosome. The HRD-TAI score is defined as the number of allelic imbalances (AIs) that extend to the telomeric ends of a chromosome without crossing its centromere. The HRD-LST score is the number of chromosomal breaks between adjacent regions of at least 10 Mb with a distance between them not larger than 3 Mb. The aggregated form of these three measures are often referred to as the HRD score. The three genomic scar scores were calculated for each sample with the help of the scarHRD R package.

### DNA methylation analysis

For determining the BRCA1 methylated cases the labeling was used from Liu et al. and to confirm the results DNA methylation data measured were downloaded from the TCGA data portal (https://portal.gdc.cancer.gov/)^11^ and the median beta values of promoter-associated probes in a given gene were used.

### Cell lines and cell culture

Human gastric cancer cell lines were obtained from the Cancer Cell Line Encyclopedia (CCLE) core facility (BROAD Institute, Cambridge), which obtained them directly from commercial sources and authenticated the lines using standard short tandem repeat analysis. RPE1 cells were obtained from ATCC. RPE1 cells were grown in DMEM-F12 (Life Technologies, #10565042) supplemented with 10% FBS and 1% penicillin/streptomycin, HGC27 cells were grown in DMEM (Life Technologies, #11965118) supplemented with 10% FBS and 1% penicillin/streptomycin. AGS, KE39, GSU, and NUGC3 were grown in RPMI 1640 (Life Technologies, #11875119) supplemented with 10% FBS and 1% penicillin/streptomycin. Cells lines were maintained in humidified 37°C the incubator with 5% CO2 and routinely tested for mycoplasma contamination (Lonza #LT07-118).

### Cell Proliferation Assay

For IC50 experiments, 1000 cells were plated in a flat-bottom 96-well plate. Cells were treated with either vehicle (DMSO) or different concentrations of Chemotherapeutic agents or PARP inhibitors. Luminescence was measured using CellTiter-Glo (Promega, #G7572) for ATP amount after 3-5 days, and final readings were normalized with Day 1 luminescence readings. For colony formation assay, 2×10^4-1×10^5 cells were plated in 6-well plates. Cells were then treated with DMSO or inhibitors, and treatments were renewed every 3-4 days (different doses of UV radiation for UV damage assay using StrataLinker 2400 irradiator). After 7-10 days, cells were fixed in 1% paraformaldehyde for 15 minutes at RT, washed twice with PBS, and stained with 0.1% crystal violet solution in ethanol (Sigma Aldrich, #HT901) for 15 minutes at room temperature. ImageJ was used to quantify the mean intensity of the scanned plates.

### *In-silico* drug sensitivity analysis

To evaluate drug sensitivity data for gastric cancer cell lines, PRISM drug repurposing data were downloaded (22Q2 data release from DepMap website of Broad Institute (https://depmap.org). The cell lines were divided into a high or low group based on quartiles of HRD score. The difference between the for the two groups was plotted as median, and the p-value was calculated using the Mann Whitney test.

### Microscopy/Image quantification

Immunofluorescence imaging was performed using Nikon Eclipse Ti2 Series inverted microscope (×40 objective). NIS-Elements AR software was used to acquire the images. DAPI channel was used to focus the cells, and the images for 488 and 567 channels were acquired, keeping the same exposure across the samples. For image analysis, the 16-bit images were converted to 2-bit images in ImageJ, followed by ‘hole filling’ and segmentation (watershed) commands. The cells were then automatically recognized and counted using ‘analyze particles’, and the file was saved as an ‘image mask’. Finally, the image mask was overlaid on the original green (488) and red (567) split channels, and the ‘measure’ command was used to get the mean fluorescence intensity from each cell.

### Single cell RNA sequencing Sample preparation

NUGC3 cells were treated with the IC50 concentration of Cisplatin, Oxaliplatin, 5-FU and DMSO for 2 days. The cells were washed with PBS, trypsinized and labeled with cell hashing antibodies, TotalSeq^™^-C0251 Hashtag 1 (BioLegend #394661) and TotalSeq^™^-C0252 Hashtag 2 (BioLegend #394662). Viable cells were washed and resuspended in PBS with 0.04% BSA at a cell concentration of 1000 cells/µL. About 17,000 viable mouse cells were loaded onto a 10× Genomics Chromium^™^ instrument (10× Genomics) according to the manufacturer’s recommendations. The scRNAseq libraries were processed using Chromium^™^ single cell 5’ library & gel bead kit (10× Genomics). Matched cell hashing libraries were prepared using single cell 5’ feature barcode library kit. Quality controls for amplified cDNA libraries, cell hashing libraries, and final sequencing libraries were performed using Bioanalyzer High Sensitivity DNA Kit (Agilent). The sequencing libraries for scRNAseq and scTCRseq were normalized to 4nM concentration and pooled using a volume ratio of 4:1. The pooled sequencing libraries were sequenced on Illumina NovaSeq S4 300 cycle platform. The sequencing parameters were: Read 1 of 150bp, Read 2 of 150bp and Index 1 of 8bp. The sequencing data were demultiplexed and aligned to mm10-3.0.0 using cell ranger version 3.1.0 pipeline (10× Genomics).

### Single cell RNA sequencing

#### General analysis

scRNA-seq IntegrateData function in Seuratv4 was used to counteract batch effects among human samples. Principal Component Analysis (PCA) was then completed on the integrated object and the quantity of principal components selected for clustering was determined using the integrated object’s elbow plot. Cells were then visualized primarily using UMAP non-linear dimensional reduction from which feature and violin plots were generated to demonstrate distribution of gene expression and expression levels of various marker genes and gene signatures throughout the population.

#### Pre-processing, alignment and gene counts

De-multiplexing, alignment to the transcriptome, and unique molecular identifier (UMI) collapsing were performed using the Cellranger toolkit provided by 10X Genomics.

#### General Clustering

Standard procedures for QC filtering, data scaling and normalization, detection of highly variable genes, and hashtag oligo (HTO) demultiplexing were followed using Seurat v4 in RStudio. Cells with unique feature counts lower than 500 and greater than 7,500 as well as cells with greater than 15% mitochondrial DNA were excluded. Counts were log-normalized and scaled by a factor of 10,000 according to the default parameters when using the Seurat LogNormalize function. Variable features were identified, and the data were scaled using the default parameters (Ngenes = 2000) of the FindVariableFeatures Extended Data Fig. ScaleData Seurat functions, respectively. HTOs were demultiplexed using the HTODemux function, and cells were identified as containing HTO-1 or HTO-2 based on their maximal HTO-ID signal. The cell population was filtered to contain only HTO-positive, singlet cells for further analysis. Principle component analysis (PCA) was completed on the remaining cells and 10 principle components were selected for clustering, tSNE, and UMAP analyses. Cells were visualized primarily using UMAP non-linear dimensional reduction (dims 1:10, resolution = 0.3), from which feature plots were generated to demonstrate distribution of gene expression and different drugs treatment cells and expression levels of various marker genes throughout the population. Marker genes for each resulting cluster were found using the FindMarkers function with the minimum prevalence set to 25%. Cluster identities were defined using known marker genes enriched in different pathways.

#### scRNA-seq gene signature analysis

To analyze existing gene signatures on our scRNA-seq data, the Seurat AddModuleScore function in Seurat v4 was used to calculate the average normalized and scaled gene expression of a given gene list in each individual cell. Specific cell types were identified using established marker genes and gene signatures. Gene signature scoring was then visualized with feature and violin plots. To generate novel gene signatures, the Seurat FindMarkers function was used to create lists of genes differentially expressed in one specified subset in comparison to another given subset. Minimum prevalence was set to 25%.

### Patients and cohorts

In this study, 791 whole genome (WGS) and whole exome sequenced (WES) pretreatment samples were analyzed from four cohorts of patients with GEA. We analyzed 68 cases of gastric cancer and 97 cases of esophageal cancer with WGS data and 441 cases of gastric cancer and 185 cases of esophageal cancer with WES data (Supplementary Table 1). TCGA and ICGC cohort**-** For the analysis, the normal, tumor bam and vcf files were downloaded from the ICGC data portal (https://dcc.icgc.org/) for the WGS, and from the TCGA data portal (https://portal.gdc.cancer.gov/) for the WES samples. The WES vcf files were generated by MuTect2 (GATK, v3.8).

### Mutation, copy number, and structural variant calling

For the WGS cohorts, germline and somatic mutations, structural variants, allele-specific copy numbers were obtained from the DCC Data Portal (https://dcc.icgc.org/) and for the TCGA WES datasets germline and somatic mutation status and the normal and tumor BAM files were downloaded via the GDC Data Portal and Sequenza^12^ was used to determine allele-specific copy number profiles. Germline and somatic mutations for all cohorts were annotated using InterVar (https://wintervar.wglab.org/) and the genotyping of the samples was performed as shown (Supplementary Figures 1-4). The samples with identified *BRCA1/2* and other HR related pathogenic gene mutations are listed in Supplementary Table 1. Further details are available in the Supplementary Methods.

### Mutational signatures

Somatic single base substitution signatures were calculated using the deconstructSigs R package (https://cran.r-project.org/web/packages/deconstructSigs/index.html), with the COSMIC signatures as a mutational process matrix (cancer.sanger.ac.uk/cosmic/signatures). Doublet base substitution and indel signatures were extracted with the ICAMS R package (https://cran.rstudio.com/web/packages/ICAMS/index.html).

Further details are provided in the Supplementary Methods.

### HRD score calculation

The scarHRD R^13^ package was used to calculate the HRD-associated genomic scar scores (HRD-LOH, HRD-LST, HRD-TAI) from the copy number profile of the samples. The sum of the HRD scores together with the extracted signatures and deletion profiles were then reported. Additional details are described in the Supplementary Methods.

### DNA methylation analysis

DNA methylation data measured by the Illumina HumanMethylation450 platform were downloaded from the TCGA data portal (https://portal.gdc.cancer.gov/)^11^. The median beta values of promoter-associated probes in a given gene were used. Additional details are described in the Supplementary Methods.

### DDB2 proteo-probe assay for NER Deficiency

The DDB2 proteo-probe assay, that detects 6,4-photoproducts was performed based on^14, 15^. Briefly, the purified HA-tagged DDB2 protein complex was used as a probe in an affinity fluorescence-based assay. Cells were plated on glass-teflon microscope slide (Tekdon, #518plain) and the next day exposed to 20 J/m2 UV-C at 254 nm using a StrataLinker 2400^™^ irradiator. 5 or 150 minutes following UV exposure, cells were fixed with ice-cold methanol for 10 minutes at room temperature. After serial rehydration with phosphate-buffered saline (PBS), non-specific binding sites were blocked by incubation with PBS-0.3% BSA. DDB2 proteo-probe was diluted in PBS-BSA and added to the fixed cells for 2 hours at 37°C. Cells were then washed twice with PBS, and rabbit anti-HA antibody (1:600; CST #3724S) was used to label the hybridized DDB2 proteo-probe. Cells were then rewashed with PBS, and goat anti-rabbit antibody coupled to Alexa fluor488 fluorochrome (1:600; Lifetech #A11008)) was added. Following two final washes in PBS and one in purified water, coverslips were mounted in Fluoro-Gel II containing DAPI (EMS #17985-51).

### RAD51 assay for HR Deficiency

Cells were plated on glass-teflon microscope slide (Tekdon, #518plain) and the next day treated with either DMSO or Phleomycin (InvivoGen # ant-ph-1) for 1 hr. Cells were then washed and fixed with 4% paraformaldehyde (PFA) for 15 min at RT, blocked and permeabilized PBS-BSA + 0.3% Triton X-100 for 15 min at RT, and incubated with primary antibodies (Rad51 Rabbit 1:200, CST # 8875 and pH2A.X Mouse 1:600 CST #80312) in PBS-BSA overnight at +4°C. After three washes with PBS, secondary antibodies; anti-Rabbit IgG Alexa Fluor 488 (Life tech #A11008) and anti-Mouse IgG Alexa Fluor 568 (Life tech #A11004), diluted 1:300 in PBS-BSA with DAPI (1:1000) for 2 hr at RT. Finally, the cells were washed with PBS three times, mounted, and covered in aluminum foil for imaging.

Additional details are described in the Supplementary Methods.

## RESULTS

### Frequency of HR deficiency in GEA whole exome sequencing cohorts

To evaluate the frequency of HR deficiency in gastrointestinal cancers, we analyzed WES and whole genome sequencing (WGS) data from patient samples. In addition to a wider variety, WGS data contains about a 100-fold greater number of HR deficiency-induced mutational events than WES data^16^. In addition, certain genomic aberrations, such as large-scale rearrangements, can be detected with high confidence only by WGS. However, significantly more cases of WES data are available for analysis compared to WGS data. (After removing MSI cases, there were 316 gastric cancer WES cases in TCGA, 68 gastric cancer WGS cases from PCAWG; there were 177 esophageal cancer WES cases in TCGA, 97 esophageal cancer WGS cases in PCAWG and 419 colorectal WES cases in TCGA versus 42 colorectal WGS cases in PCAWG; Supplementary Table 1). We therefore began our analysis using WES cohorts.

Mutations in the canonical HR genes *BRCA1, BRCA2* and *PALB2* are present in 3-5% of GEA tumors^17^; however, only a fraction of these cases have both a predicted loss-of-function mutation as well as loss of heterozygosity (LOH) of the wild-type (WT) allele (Supplementary Figures 1-4). Since LOH of the WT allele is typically required to confer HR deficiency^18^ in the setting of a *BRCA1/2* mutation, we considered only those *BRCA1/2* mutant cases that had biallelic mutations or a mutation accompanied by LOH to be HR deficient.

In the TCGA gastric cancer WES dataset, we identified two cases with a pathogenic *BRCA2* mutation accompanied by LOH, one BRCA2 case where both alleles had pathogenic mutations, and two cases with pathogenic *BRCA1* mutation accompanied by LOH (Figure 1, Supplementary Table 2). We also found four pathogenic PALB2 mutations accompanied by either LOH or biallelic mutations in the TCGA gastric cancer cohort (Figure 1, Supplementary Table 2). In the TCGA esophageal and colorectal WES data sets, we identified one case in each cohort with a pathogenic *BRCA2* mutation accompanied by LOH.

**Figure 1.**
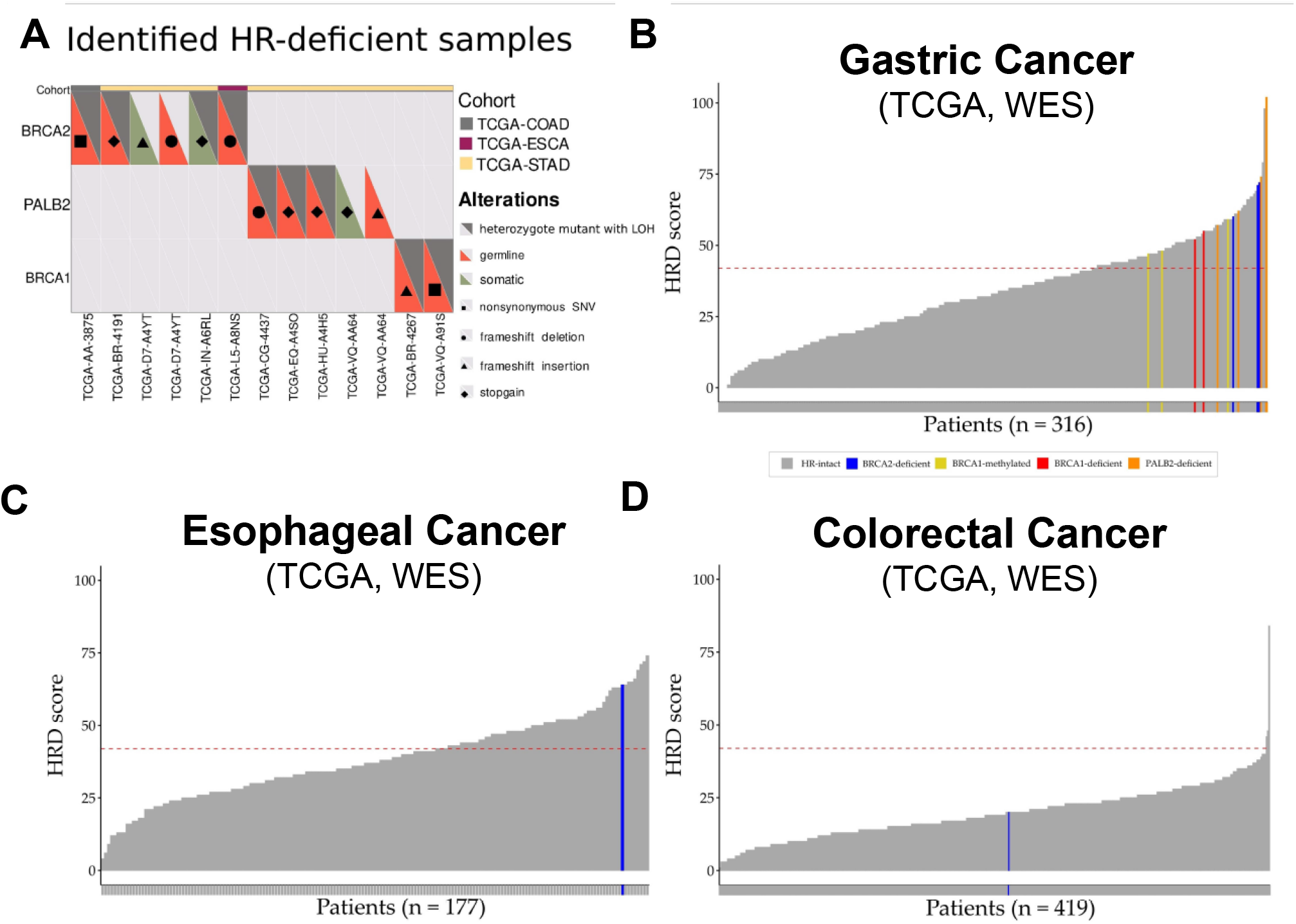
HRD mutational signature in gastrointestinal malignancies. A. Germline and somatic mutations and copy number alterations of HR genes across the TCGA-STAD, TCGA-ESCA, TCGA-COAD and TCGA-READ WES cohorts. B. -D. HRD score distribution in the TCGA-STAD (B), TCGA-ESCA (C), TCGA-COAD and TCGA/READ (D) WES samples. Cut-off value of ≥42 for HR deficiency was previously defined (dashed-line).

We next determined mutation and LOH status for DNA damage checkpoint genes such as *ATM, ATR, CHK2, TP53*, and *RB1* (Supplementary Figures 1-4), although loss of function of these genes is usually not associated with an HR deficiency mutational signature^19^. *TP53* was the most frequently altered DNA damage checkpoint genes in the cohorts. We also identified three gastric adenocarcinoma cases with significant BRCA1 promoter methylation (Figure 1). These analyses indicate that a small but significant fraction of gastrointestinal cancers harbor mutations in genes that function in DNA repair and DNA damage response.

### Genomic scar-based HRD scores in gastrointestinal cancer

The first FDA approved genomics-based method to quantify the degree of HRD was derived based on data obtained from hybridization microarrays. It has three components: (1) The HRD-LOH score is calculated by tallying the number of LOH regions exceeding 15 Mb in size but less than the whole chromosome^20^ ; (2) the Large-scale State Transitions (LST) score^21^ is defined as the number of chromosomal breaks between adjacent regions of at least 10 Mb, with a distance between them not larger than 3Mb; and (3) the number of Telomeric Allelic Imbalances (TAI)^22^ is the number of AIs (unequal contribution of parental allele sequences) that extend to the telomeric end of a chromosome. These measures were later adapted to next generation sequencing and the sum of these scores is referred to as the HRD score^8^, which was recently approved by FDA as a companion diagnostic for prioritizing patients with ovarian cancer for PARP inhibitor therapy.

In the TCGA WES gastric cancer cohort, three *BRCA2* deficient, four *PALB2* deficient, two *BRCA1* deficient and three *BRCA1* promoter methylated cases all had an HRD score ≥42 (Figure 1*A*, p-value < 1×10^−5^, Fisher exact test), which is the threshold for HR deficiency previously defined for ovarian cancer^8^. However, there were an additional 89 of 316 (28%) cases without a *BRCA1/2* or other HR gene mutation that also had an HRD score ≥42 (Figure 1*B)*. Out of these 89 samples 4 could be explained by TOP2A mutation associated genomic instability^23^ (Supplementary Figure 5). Of the three molecular subtypes of gastric adenocarcinoma included in our analysis (MSI cases were excluded)^3, 4^, substantially more cases with an HRD score ≥42 belonged to the subgroup with chromosomal instability CIN, in contrast to the genomically stable and EBV positive subtypes (Supplementary Figure 6).

In the TCGA WES esophageal cancer cohort, the *BRCA2* deficient case also demonstrated an HRD score ≥42 (Figure 1*C*). Similar to the TCGA gastric cancer cohort, there were an additional 68 of 179 cases (40%) without a *BRCA1/2* or other HR gene mutation that also had an HRD score larger than the threshold. Given the high frequency of HR deficiency in GEA, we analyzed the prevalence of HR deficiency in another common gastrointestinal malignancy, colorectal cancer. However, unlike GEA, HR deficiency was rare in colorectal cancer. In a cohort of 419 colorectal cancer, only 3 had an HRD score ≥42, suggesting that HRD is rare in this cancer type (Figure 1*D)*.

Determination of HRD scores from WES data can be impacted by the relatively low number of SNVs (24) ; therefore, we also calculated HRD scores from WGS data for the 32 cases that had both WGS and WES data available in the TCGA STAD cohort. Similar to other solid tumors, the WES and WGS derived HRD scores showed a strong correlation (*R* = 0.87; p = 1.4×10^−10^, Supplementary Figure 7).

Since only a minority of cases with high levels of HR deficiency associated mutational signatures harbored a loss of function in *BRCA1/2* or other HR genes, we investigated whether the detected HR deficiency could be explained by other mechanisms. Suppression of HR gene function can occur via other mechanisms such as promoter methylation of HR genes, such as *BRCA1* in breast cancer, or regulatory proteins such as *RBBP8*^24-26^. However, in our analyses, we were unable to find evidence of promoter methylation that could explain the increased HRD scores, and other HR deficiency associated mutational signatures (Supplementary Figure 8). We found no other HR related genes whose promoter methylation was associated with increased HRD scores in the analyzed cohorts.

### High HRD score by mutation signature analysis correlates with platinum and PARP inhibitor sensitivity in gastric cancer cell lines

To evaluate whether putative HR deficient gastric cancer cases, as identified by mutational signatures, are in fact HR deficient, we calculated the HRD score for 31 gastric cancer cell lines and stratified them by increasing HRD score. An HRD score cutoff of 42 was used to divide cell lines into high and low categories for further investigation (Figure 2*A*). It is worth noting, similar to patient data (Figure 1), high HRD score cells lines do not necessarily harbor alterations in BRCA1/2 or other categories of HR-related genes (Supplementary Figure 9). Next, to test whether gastric cancer cell lines with high HRD scores are functionally HR deficient, we employed a RAD51 functional assay which assesses HR proficiency by interrogating the ability of RAD51 foci to form in response to double strand DNA (dsDNA) breaks. We chose two high HRD score (KE39 and HGC27) and two low HRD score (AGS and GSU) gastric cancer cell lines to perform the assay (Figure 2*A*). In addition to these four gastric cell lines, we utilized RPE1 as a non-neoplastic, HR proficient cell line control for our further experiments. After treating the cells with radiomimetic DNA-cleaving agent phleomycin for one hour, phospho-H2AX (pH2AX) and RAD51 were detected by immunofluorescence, indicating the presence of dsDNA breaks and HR activity, respectively. Irrespective of HRD scores, three of the cell lines displayed robust pH2AX activity in response to phleomycin treatment and could be further evaluated for the development of RAD51 foci; RPE1 and GSU cells did not display DNA damage in response to a standard phleomycin dose, suggesting either faster repair or insufficient DNA damage (Supplementary Figure 10*A-B*). While the low HRD score cell line AGS induced RAD51 foci in response to DNA damage, indicating functional HR, both high HRD score cell lines did not display RAD51 foci, confirming HR deficiency (Figures 2*B-C*). These results validate that high HRD scores observed by mutation signature analysis can identify gastric cancer cell lines with functional HR pathway deficiency.

**Figure 2.**
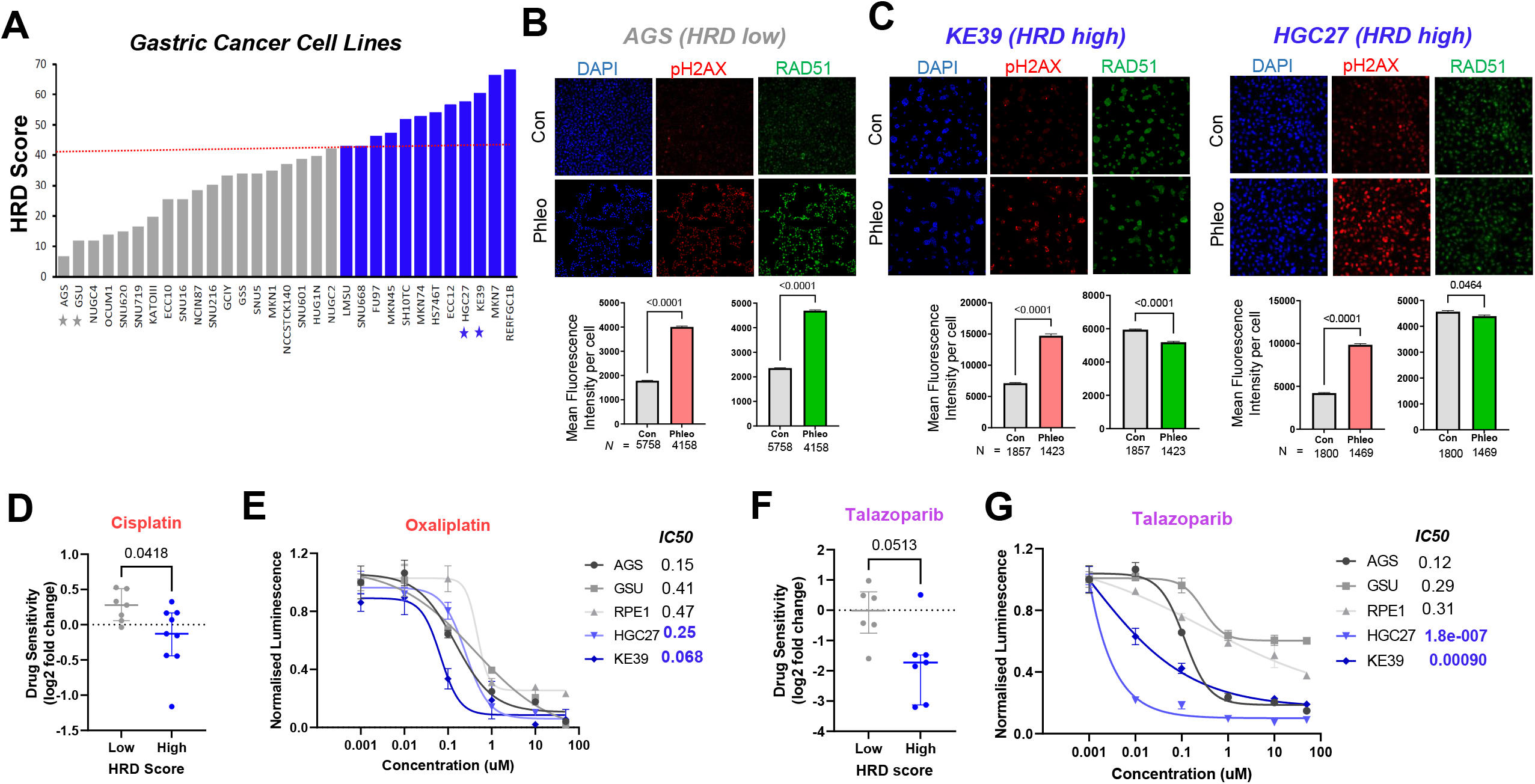
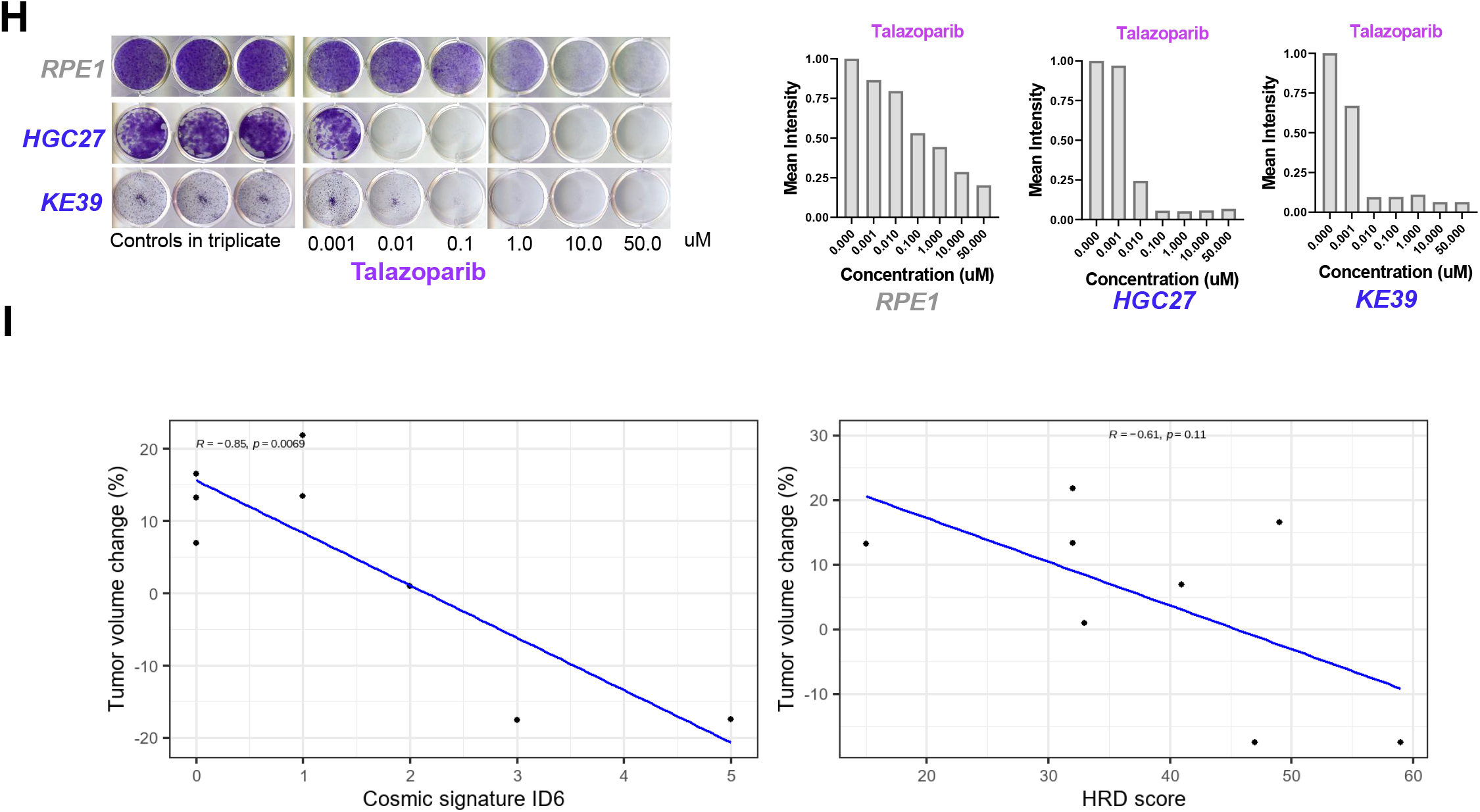
High HRD scores by mutation signature analysis is associated with platinum and PARP inhibitor sensitivity in gastric cancer. A. Gastric cancer patients cell lines from CCLE stratified by HRD Score. Cut-off value of ≥42 for HR deficiency was previously defined (dashed-line); cell-lines with low (gray) and high (blue) HRD scores indicated. Starred cell-lines used in experimental validation studies. B. Representative immunofluorescence images (left) and quantification (right) for AGS cells; pH2AX (red) indicating ds-DNA breaks and RAD51 (green) indicating DNA repair after treatment with DMSO control or radiomemetic (gamma-ray) phleomycin. C. Representative immunofluorescence images (left) and quantification (right) from KE39 and HGC27 cell lines; pH2AX (red) and RAD51 (green) indicating DNA repair after treatment with DMSO control or radiomemetic (gamma-ray) phleomycin. D. Cisplatin sensitivity between low (gray) and high (blue) HRD gastric cancer cell lines available in BROAD institute PRISM repurposing drug screen dataset. Difference between the cisplatin sensitivity (log2 fold change) is represented as the median. E. Dose-response curve of non-neoplastic RPE1, low (gray) and high (blue) HRD (blue) gastric cancer cell lines to oxaliplatin. Best-fit IC50 scores are displayed. F. Talazoparib sensitivity between low (gray) and high (blue) HRD gastric cancer cell lines available in BROAD institute PRISM repurposing drug screen dataset. Difference between the cisplatin sensitivity (log2 fold change) is represented as the median. G. Dose-response curve of non-neoplastic cell line RPE1, low (gray) and high (blue) HRD gastric cancer cell lines to indicated concentrations of Talazoparib. Best-fit IC50 scores are displayed. H. Colony formation assay (left) and the quantification shown as mean intensity from each well using ImageJ (right) of gastric cancer cell with high HRD score (blue) and non-neoplastic RPE1 (gray). I. Correlation between tumor volume change (%) and Cosmic signature ID6 absolute numbers and HRD score. Pearson correlation coefficient (−0.85 and −0.61) and p-values (0.0069 and 0.11) were calculated. *All data expressed as mean ± SD and P-values calculated by Mann Whitney test unless otherwise indicated

Our and others’ prior work have shown that HR-deficient ovarian, breast, and prostate cancers are exquisitely sensitive to platinum agents and PARP inhibition^9, 16, 25-27^, and we hypothesized that gastric cancers deficient in HR (or with high HRD scores) would be more sensitive to platinum agents and PARP inhibition. To test this hypothesis, we first used the chemotherapeutic agents’ sensitivity data from the Broad Institute’s PRISM repurposing drug screen^28^. Gastric cancer cell lines were divided into two groups/quartiles based on low and high HRD scores (Figure 2*A*). The sensitivity to most chemotherapeutic agents is not significantly different between these two groups (Supplementary Figure 11). Indeed, gastric cancer cell lines evaluated in this study did not show differential sensitivity to 5-FU treatment (Supplementary Figure 12*A*). Although most chemotherapy agents did not display a difference in killing activity, platinum chemotherapy did show differential activity in low and high HRD scoring cell lines. High HRD score cell lines showed greater sensitivity to cisplatin compared to low HRD score cell lines (Figure 2*D*), which we confirmed for HGC27 but not KE39 gastric cancer cell lines (Supplementary Figure 12*B*). Closer examination of Oxaliplatin treated cell lines showed a subset of high HRD score cell lines with greater sensitivity (Supplementary Figure 11, red box). Consistently, luminescence-based viability assays indicated that HR-deficient cell lines KE39 and HGC27 were predominantly more sensitive to oxaliplatin (Figure 2*E*). These results suggest that HR-deficient gastric cancer display greater sensitivity to platinum agents.

We next investigated whether HR-deficient gastric cancer cell lines are sensitive to PARP inhibition. Humans have at least three important PARP family members (PARP1/2/3), and each PARP inhibitor has different potency, with talazoparib (BMN-673) exhibiting the most potency^29-31^. Using Broad PRISM drug sensitivity data, we found that high HRD score gastric cancer cell lines showed greater sensitivity to many PARP inhibitors compared to low HRD score lines (Supplementary Figure 13), with talazoparib showing the most potent effect (Figure 2*F*). We confirmed that high HRD score lines showed exquisite sensitivity to talazoparib compared to low HRD score lines by cell-titer go cytotoxicity assays (Figure 2*G*). Olaparib, rucaparib, AZD2461, and velaparib also showed greater activity in high HRD score cell lines (Supplementary Figure 12*C-F*). Since the duration of PARP inhibition is important^32^, we also performed two-week colony formation assays with talazoparib and veliparib, showing greater sensitivity in high HRD cell lines (Figure 2*H* and Supplementary Figure 12*G*).

### HRD associated mutational signatures are associated with response to PARP inhibitor treatment in the clinical setting

In a recent clinical trial (NCT03008278), patients with advanced GEA were treated with a combination of olaparib and ramucirumab, with several patients showing significant response. Of the 49 patients enrolled, 9 cases have been profiled by whole exome sequencing so far. We calculated the various HRD associated mutational signature scores for these patients and found either significant association (ID6) or strong tendency (HRD score) for association between benefit from olaparib containing therapy and HRD associated mutational signatures (Figure 2*I*). While this analysis needs to be extended for the entire cohort and further validated, it suggests that HR deficient GEA cases identified by mutational signatures may significantly benefit from PARP inhibitor-based therapy.

### NER deficient gastric cancer cell line displays cisplatin and PARP inhibitor sensitivity

Based on the experimental confirmation that HR-deficient gastric cancers are sensitive to platinum chemotherapy, we wondered whether HR-deficient cell lines could be identified by sensitivity to cisplatin. To this end, we analyzed Sanger’s Genomics of Drug Sensitivity in Cancer Project (GDSC)^33^, searching for gastric cancer cell lines with exquisite cisplatin sensitivity. NUGC3 gastric cancer cell line displayed the highest sensitivity to cisplatin by a significant margin compared to other gastric cancer cell lines (Figure 3*A*). Luminescent-based viability assay and colony formation assay confirmed this high sensitivity of NUGC3 cells to cisplatin (Figure 3*B* and Supplementary Figure 14*A*). Compared with cisplatin, oxaliplatin and 5-FU were not as active in NUGC3 cells by luminescent-based viability assays (Supplementary Figure 14*B-D*). These findings were consistent with the GDSC data (Supplementary Figure 14*E*).

**Figure 3.**
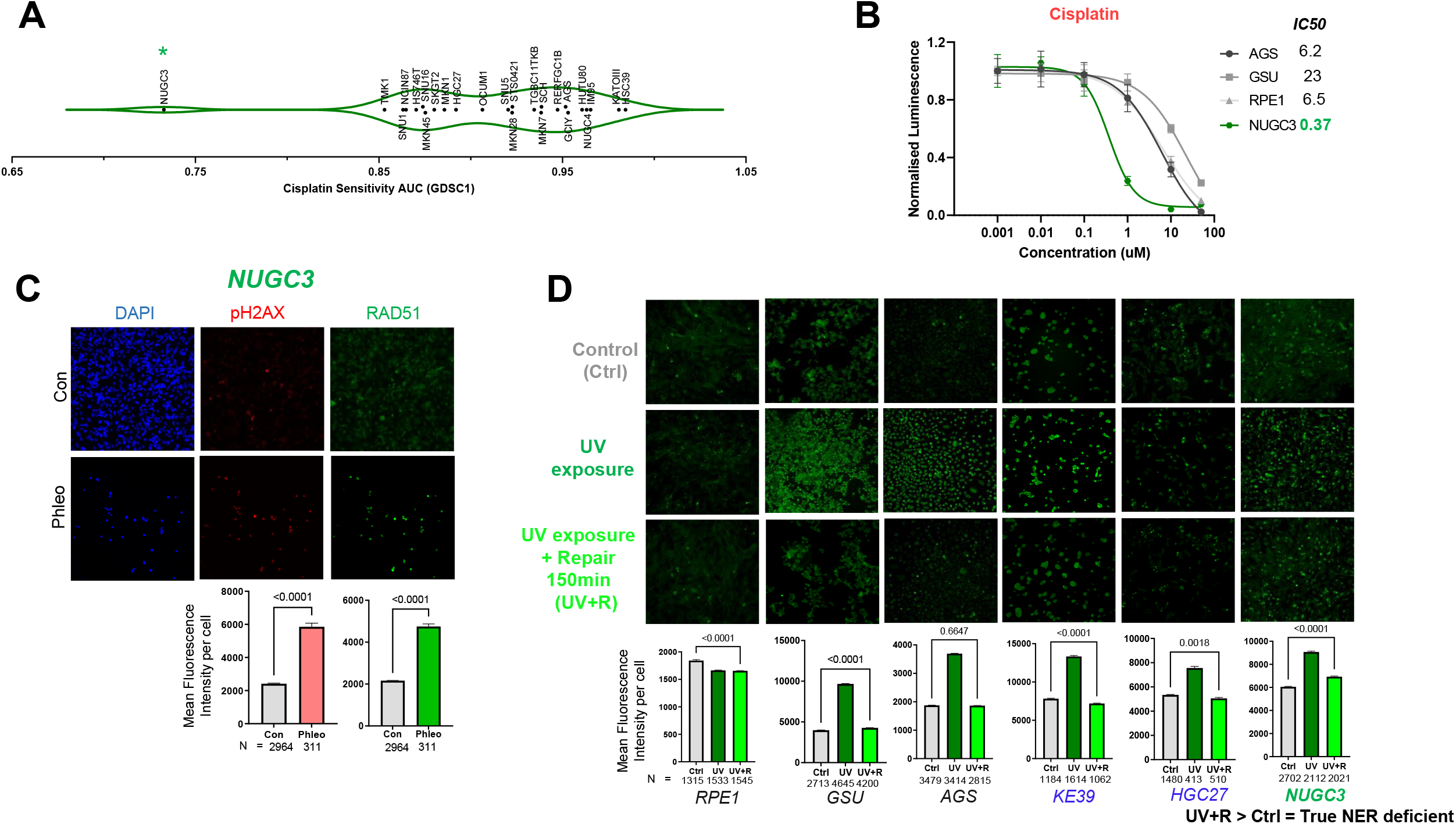
Cisplatin sensitivity analysis reveals NER deficient gastric cancer cell line. A. Cisplatin sensitivity in gastric cancer cell lines from Sanger’s Genomics of Drug Sensitivity in Cancer Project (GDSC1); drug response measured as Area Under the Curve (AUC); NUGC3 (green star). B. Cisplatin dose-response curve of NUGC3 (green) and three control cell lines (gray). Best-fit IC50 scores are displayed. C. Representative immunofluorescence images (left) and quantification (right) of pH2AX (red) and RAD51 (green) in NUGC3 cells after treatment with radiomemetic (gamma ray) phleomycin. D. Representative immunofluorescence images (top) and quantification (bottom) for the DDB2 proteo-probe assay to detect GG-NER defects. DDB2 signal (green) indicates DNA damage after UV treatment (dark green bars). In cells with intact GG-NER, DDB2 signal reduces to the signal in the control cells (grey bars) by 150 minutes (light green bars), indicating repair. In cells lacking NER, the signal does not completely reduce to that in control cells (UV+R > Ctrl = True NER deficient); control (gray), HR-deficient (blue), and NER-deficient (green) cell lines. *All data expressed as mean ± SD and P-values calculated by Mann Whitney test unless otherwise indicated

We next evaluated whether NUGC3 is functionally HR deficient by assessing the RAD51 assay. Phleomycin treatment yielded robust induction of pH2AX that corresponded to RAD51 foci induction, suggesting that NUGC3 is HR proficient (Figure 3*C*). We, therefore, asked whether another DNA repair pathway was defective in NUGC3 cells to explain the outlier sensitivity to Cisplatin. Nucleotide excision repair (NER) is a critical DNA repair pathway involved in the repair of UV radiation-induced DNA damage and cisplatin-induced DNA damage^15, 34^. Inactivation of this pathway, through deleterious ERCC2 mutations, is associated with cisplatin sensitivity in bladder cancer^35^. To investigate whether NUGC3 cells are NER deficient, we employed a recently developed assay that relies on specific binding of a DDB2 (damaged-DNA binding protein 2) proteo-probe to UV-induced 6,4-photoproducts. DDB2 is an essential initiation factor involved in the global-genome repair (GGR) NER pathway^36^. Following UV exposure, NER proficient lines will repair 6,4-photoproducts after 150 minutes, which is detected by a DDB2 proteo-probe hybridization returning to baseline levels. By contrast, NER deficient lines will retain UV-induced 6,4-photoproducts and the proteo-probe signal will not return to baseline. This assay demonstrated that MDA-MB-468 breast cancer cells are NER deficient^14, 15^, which corresponded to its sensitivity to cisplatin and PARP inhibitors. We applied this assay to the six lines used in our study. Similar to phleomycin treatment, RPE1 cells appeared to be resistant to UV-induced DNA damage (Figure 3*D*). All gastric cancer cell lines developed 6,4-photoproducts immediately following UV exposure as measured by DDB2 immunofluorescence (Figure 3*D*). Unlike the two HR-proficient and two HR-deficient cell lines, NUGC3 cells did not fully repair 6,4-photoproducts as indicated by retained DDB2 signal 150 min post UV exposure, indicating that this cell line carries functional deficiencies in NER. Since NUGC3 cells were unable to completely repair UV-induced 6,4-photoproducts, we next asked whether NUGC3 cells are selectively sensitive to UV radiation. Compared to RPE1 cells, NUGC3 cells displayed a dose-dependent sensitivity to UV radiation exposure by colony formation assay (Supplementary Figure 15*A*). These findings suggests that a subset of gastric cancers could be NER deficient and might respond to cisplatin-containing regimens.

To provide a potential molecular explanation for NER deficiency in NUGC3 cells, we analyzed mutation profile of these cell lines. Consistent with HR proficiency in NUGC3 cells (Figure 3*C*), we did not observe mutations in common HR genes like *BRCA1/2, PALB2, ATM, FANCA*. By contrast, we observed nonsynonymous mutations in genes that function in both arms of the NER pathway: global genome (GG-NER) and transcription-coupled (TC-NER)^34^ (Supplementary Figure 15*B*). Of all surveyed gastric cancer cell lines in the CCLE, NUGC3 is the only gastric cancer cell line harboring mutations in essential NER pathway genes (Supplementary Figure 15*C*). NUGC3 cells also showed the lowest expression of *XPC*, a gene that plays an essential role during the first step of GG-NER (Supplementary Figure 15*D*). Interestingly, lower expression of XPC correlates with better cisplatin sensitivity in gastric, ovarian, colorectal, and lung cancer^37-41^. It is worth mentioning that the only other NER-deficient cancer cell line (MDA-MB-468) reported previously, does not harbor mutations in NER genes; instead, NER deficiency in this cell line is driven by epigenetic silencing of NER gene *ERCC4*^15^. These results indicate that somatic mutations in essential NER pathway genes can contribute to functional NER deficiency in gastric cancer.

PARP is not only a vital component of the HR but also the NER pathway^42^. Several studies suggest efficacy of PARP inhibitors for ovarian and lung cancers harboring mutations in NER pathway genes (*ERCC1, ERCC8, DDB1, XAB2*)^43-45^. We, therefore, examined whether NER-deficient gastric cancer is sensitive to PARP inhibition using NUGC3 as our model. While most PARP inhibitors showed modest activity, Talazoparib showed potent, selective killing in NUGC3 cells compared to three control cell lines (Supplementary Figure 16). These data suggest there may be a role for potent PARP inhibitors, such as talazoparib, in NER-deficient gastric cancer cases.

### Identification of NER deficient gastrointestinal cancer cases in the clinical setting

Despite the potential clinical actionability of tumor NER deficiency, there are currently no functional or IHC assays available to reliably identify NER deficiency in clinical specimens. Next-generation sequencing (NGS)-based mutational signatures can be used to identify NER deficiency. We recently identified a complex mutational signature, predominantly driven by the mutational signature SBS5 and ID8, that is significantly increased in NER deficient bladder cancer cases, especially those with ERCC2 mutations^10^. We determined the values of the same complex mutational signature for the TCGA GEA cancer cases. When we used the same threshold value (>0.7) that distinguished NER deficient and proficient cases in bladder cancer, we found 22 gastric cancer cases, 14 esophageal cancer cases and 25 CRC cases with a >0.7 value (Figure 4). However, none of these cases had an inactivating mutation in the canonical NER genes (such as ERCC2 and ERCC3). In order to determine whether this complex mutational signature may in fact reflect NER deficiency we turned to the genomic analysis of eight gastric adenocarcinoma organoids derived from patients at Dana-Farber Cancer Institute from the Cancer Cell Line Factory (CCLF). One of the tumors had inactivating mutations in ERCC2 and ERCC6, two key genes of NER. The NER associated mutational signatures were the highest in this case, relative to other cases without inactivating mutations in NER genes (Supplementary Figure 17). This was a 73-year-old man who had a locally advanced, poorly differentiated, microsatellite stable gastric adenocarcinoma. He had 2 months of 5-fluorouracil (5-FU) and oxaliplatin (FOLFOX) then underwent a total gastrectomy with D2 lymph node dissection. Pathology showed minimal response to neoadjuvant FOLFOX and the pathologic stage was ypT4N0. He then switched to docetaxel, cisplatin and 5-FUfor a total of 4 cycles of adjuvant therapy. He is still alive today with no evidence of disease at the last imaging follow up, which was 4.5 years after completion of adjuvant chemotherapy. Consistent with our cell line data, this tumor was resistant to oxaliplatin-containing therapy, and perhaps the switch to cisplatin (in addition to the standard-of-care tumor resection) contributed to his excellent outcome. These data suggest patients with gastrointestinal cancer may have tumors with NER deficiency that can be detected by mutation signature analysis. Interestingly, for the same organoid case the calculated HRD score components were the lowest in the analyzed samples.

**Figure 4.**
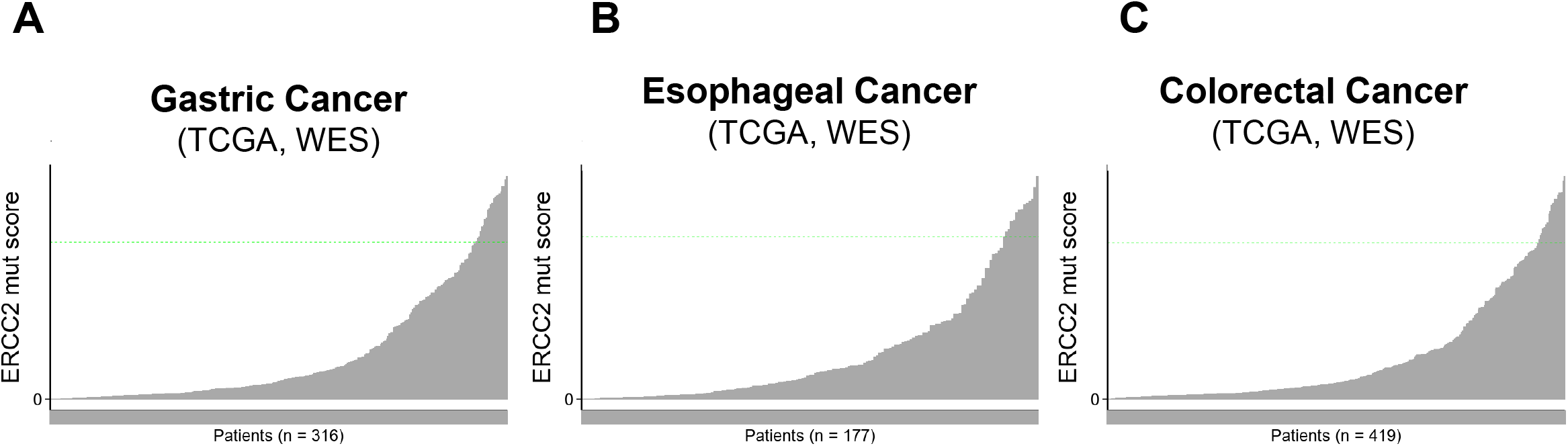
NER deficiency associated mutational signature score in gastrointestinal malignancies. The NER deficiency associated complex mutational signature was calculated in the WES data for gastric (panel A), esophageal (panel B) and colorectal (panel C) cancer WES data. The cut-off value was set to ≥0.7 for NER deficiency.

### scRNA-sequencing reveals ferroptosis as a cisplatin-specific mechanism of cell death in NER deficient gastric cancer

To better understand the sensitivity of NER-deficient gastric cancer to cisplatin, we performed single-cell RNA sequencing (scRNA-seq) on NUGC3 cells treated with sublethal doses of three different chemotherapeutics commonly used to treat gastric cancer: cisplatin, oxaliplatin or 5-FU (Figure 5*A*, Supplementary Figure 18*A*). UMAP representation of single-cell gene expression profiles demonstrated six distinct clusters categorized based on gene ontology (Figure 5*A*, Supplementary Figure 18, Supplementary Table 3 and 4). Cluster 0 (red) was induced in all three chemotherapy treatment groups (1.4% in DMSO; 44.7%, 42.1%, and 53.6% in cisplatin, oxaliplatin, and 5-FU treated, respectively); it showed a gene expression profile most significantly associated with apoptosis (Figure 5*A-C* and Supplementary Figure 18*B*), which is consistent with a general effect of chemotherapy irrespective of NER. In particular, it appears that apoptosis was induced by TNFα/NF-κB and p53 signaling (Supplementary Figure 18*C-E*).

**Figure 5:**
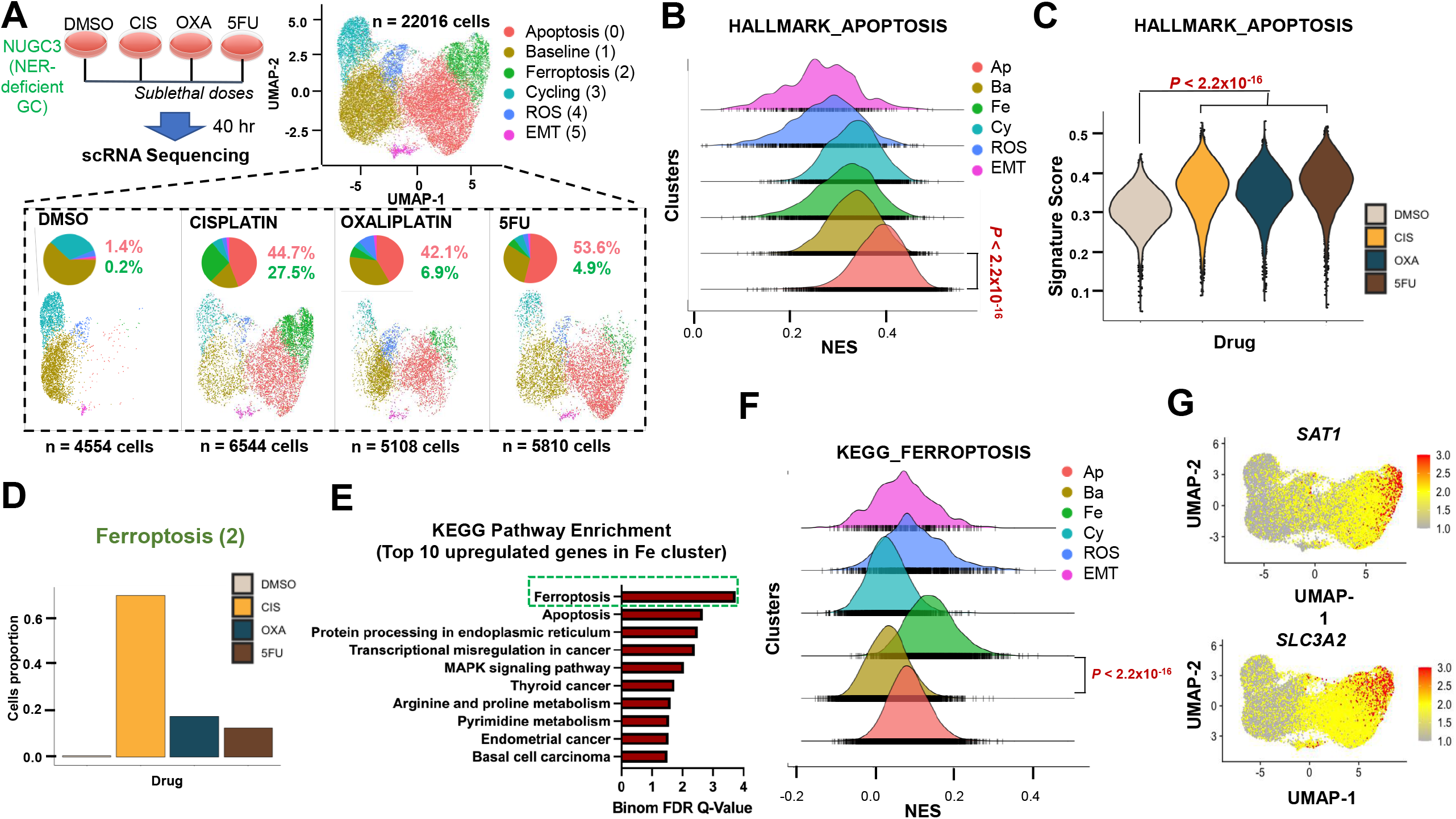
scRNA-seq analysis of NER deficient gastric cancer cells reveal distinguishing features of distinct chemotherapy treatments. A. Experimental setup and UMAP representation of single cell transcriptome profiling of NUGC3 cells treated with DMSO (control), cisplatin, oxaliplatin or 5-FU colored by distinct cell clusters. UMAP of separated samples along with pie chart indicating distribution of cell clusters (bottom panel). B. Single cell-gene set enrichment analysis (SC-GSEA) between apoptosis (0) cluster and rest of the clusters showing enrichment of HALLMARK_APOPTOSIS Pathway; NES: Normalized Enrichment Score. C. Violin plots indicating expression of HALLMARK_APOPTOSIS pathway in the four treatment groups (EnrichR) of top 20 genes upregulated in cluster Apoptosis (0) (also Supplementary table 3). P-value calculated by Wilcoxon rank sum test with Bonferroni correction. D. Normalized cell counts in each treatment group for Ferroptosis (2) cluster. E. Hallmark pathways enrichment analysis (EnrichR) of top 20 genes upregulated in Ferroptosis (2) cluster. F. Single cell-Gene set enrichment analysis (SC-GSEA) between Ferroptosis (2) cluster and rest of the clusters showing enrichment of KEGG_FERROPTOSIS Pathway; NES: Normalized Enrichment Score. G. UMAP representation of SAT1 (top) and SLC3A2 (bottom) gene expression.

By contrast, cluster 2 (green) was preferentially induced in cisplatin-treated NUGC3 cells relative to the other chemotherapeutic agents (Figure 5*D*; 27.5% in cisplatin compared to 6.9% in oxaliplatin and 4.9% in 5-FU). Pathway enrichment analysis indicated that ferroptosis was the top pathway enriched in cluster 2 (Figure 5*E-F*). Indeed, *SAT1* and *SLC3A2*, two well described ferroptosis genes^46-48^, were preferentially induced in cluster 2 (Figure 5*G* and Supplementary Figure 18*F*). These results suggest that while general apoptosis is induced by all three chemotherapeutic agents, cisplatin treatment induces an additional mechanism of cell death through ferroptosis, potentially explaining the enhanced sensitivity of NUGC3 cells to cisplatin.

We next inquired the status of TCR- and GG-NER genes, expecting that most would be silenced by way of genomic alteration (Supplementary Figure 15*C*). Interestingly, expression of POLR gene family members, a principal component of the TCR-NER pathway, was upregulated in cluster 2 (green), with *POLR2L* showing the highest expression (Supplementary Figure 18*H-I*). These results suggest that since GG-NER is compromised due to *XPC* deficiency, TCR-NER pathway components may be upregulated in a compensatory role attempting to facilitate NER. However, apart from POLR2L, which is unaltered and presumed to be functional in NUGC3 cells, the genes in the shared-NER pathway (*POLE* and *RPA1)* and downstream of TCR-NER pathway (*ERCC6* and *GTF2H1*) remain inactivated leading to functional NER deficiency (Supplementary Figure 15*B* and Supplementary Figure 18*G*), especially in response to cisplatin (cluster 2). These results implicate ferroptosis as an additional mechanism of cell death in NER deficient gastric cancer treated with cisplatin.

## DISCUSSION

Homologous recombination deficiency is often driven by loss of function of *BRCA1*/*BRCA2* or *PALB2* in tumor cells, which is frequently the result of an inactivating mutation coupled with the loss of the accompanying wild type allele (loss of heterozygosity). Consequently, many HR deficient cases are identified by sequencing *BRCA1/2 or PALB2*, and this approach has led to initial approval of PARP inhibitors in multiple tumor types including breast, ovarian, and prostate cancer. It is clear that mechanisms beyond *BRCA1/2* or *PALB2* loss may also drive tumor HR deficiency. These mechanisms include loss or mutation of other HR genes (such as *RAD51C*, and others), suppression of expression of *BRCA1* or other HR genes by methylation, and probably through other yet-to-be characterized mechanisms.

Mutational signature-based approaches have recently been applied to improve prediction of HR deficiency because they detect the consequences of HR deficiency rather than the underlying cause. The first diagnostic HR deficiency mutational signature (HRD score) was recently approved to direct PARP inhibitor therapy in ovarian cancer cases without *BRCA1/2* mutations based on data showing that patients with high HR deficiency associated mutational signatures without canonical HR gene mutations (*BRCA1, BRCA2*, etc.) benefited from PARP inhibitor therapy^9^. These findings raise the possibility that other tumors with similar mutational signatures in the absence of core HRD mutations may also be functionally HR deficient and could benefit from PARP inhibitor therapy.

GEA are only partially characterized in terms of DNA repair pathway aberrations. We find that *BRCA1/2* loss-of-function mutations – when present in both alleles or when coupled with LOH – are associated with the same HR deficiency associated mutational signatures as are present in *BRCA1/2*-mutant ovarian, breast, or prostate tumors. Therefore, *BRCA1/2* mutant GEA, although rare, appear to have *bona fide* HR deficiency.

In addition to the small percentage of HR-deficient upper gastrointestinal cancers due to loss of *BRCA1/2*, we also found that a significant number of GEAs with WT *BRCA1/2* showed levels of HRD associated mutational signatures as high as those observed in *BRCA1/2* deficient cases. Reliable detection of HR-deficient GEA may have important clinical implications. Patients with HR-deficient tumors may be prioritized for platinum-based chemotherapy^9, 49^.

HR deficiency associated mutational signatures were first identified in breast and ovarian cancer, which harbor the highest frequency of *BRCA1/2* inactivating events. We now show that the same signatures may also be useful in identifying HR deficiency in GEA, a tumor type with far less frequent alterations in *BRCA1/2*. These findings may have implications for PARP inhibitor clinical trials and suggest that mutational analysis of known HR genes such as *BRCA1/2* could be combined with mutational signature approaches (such as the HRD score) to identify cases most likely to harbor functional HR deficiency. Additional studies will be required to define mechanisms of HR deficiency in GEA, optimize threshold values of the HR deficiency mutational signatures for clinical application, and understand the therapeutic implications of HR deficiency in GEA. We further validated that three FDA approved PARP inhibitors, along with Veliparib and AZD2461, which are in preclinical and clinical trials, showed better sensitivity and tolerability than the classic PARP inhibitors in some of our studies ^32, 50^.

NER deficient cell line NUGC3 has loss of function mutations in the genes that participate in GG (*XPC, CHD1L, MCR51*) and TCR NER pathways. While XPC is necessary for the first repair step of GG-NER, RNA polymerase II encoded by the POLR2L gene is necessary for the first repair step of TCR-NER^51^. NUGC3 cells have loss of function mutation in *XPC*, but *POLR2L* is wild type though other downstream genes of *POLR2L* involved in subsequent TCR-NER repair steps have loss of function mutations, e.g., *ERCC6 and GTF2H1*^*52*^. The shared-NER pathway also carries loss of function mutations (*POLE and RPA1*). Interestingly, The *CHD1L* mutation found in NUGC3 cell line is a pathogenic (score 0.91) breast cancer somatic mutation (Genomic Mutation ID COSV63615250).

Our study has some limitations. 1) As opposed to WGS, the limitations of using WES samples are fewer available variants for the mutational signature extraction part, decreasing the reliability of the extracted features. 2) The extracted mutational signatures are limited to those previously described within TCGA-STAD, TCGA-ESCA, TCGA-COAD, and TCGA-READ cohorts, which raises the possibility of missing novel signatures. 3) The extracted mutational signatures and genomic scar scores only inform about the historical state of the tumors, and do not engender information about possible acquired drug resistance mechanisms. 4) We used six gastric cancer cell lines to validate HR/NER deficiency and drug sensitivity. The validation can be scaled up to a high-throughput fashion using automated imaging, incorporating more GEA cell lines in future studies. Furthermore, futures studies should incorporate newer models, such as organoids. 5) We did not combine chemotherapeutic agents and PARP inhibitors in our studies; combinations may show better sensitivity than monotherapy but may manifest unnecessary toxicity. 6) Lastly, our results need confirmation from the perspective of clinical trials to show that the patients with GEA that harbor deficiency in HR and NER pathways are more responsive to platinum-based chemotherapy and selective PARP inhibitors.

## ACKNOWLEDGMENTS

Results shown here are based in part from data generated by the TCGA Research Network: http://cancergenome.nih.gov/ and the International Cancer Genome Consortium (ICGC): https://icgc.org/. The results presented in the current publication are based in part on the use of study data downloaded from the dbGaP web site.

## Abbreviations

5-FU: Fluorouracil
CCLE: Cancer Cell Line Encyclopedia
COAD: Colon adenocarcinoma
DDB2: DNA damage-binding protein 2
ESCA: Esophageal carcinoma
GDC: Genomic Data Commons
GEA: Gastric and esophageal adenocarcinoma
GG-NER: Global genome-nucleotide excision repair
HR: Homologous recombination
LOH: Loss of heterogeneity
LST: Large-scale State Transitions
NER: Nucleotide excision repair
PARP: Poly (ADP-ribose) polymerase
scRNA-seq: Single cell RNA sequencing
STAD: Stomach Adenocarcinoma
TAI: Telomeric Allelic Imbalances
TCGA: The Cancer Genome Atlas
TC-NER: Transcription coupled-nucleotide excision repair
VCF: Variant Call Format
WES: Whole exome sequenced
WGS: Whole genome sequencing
XPC: XPC Complex Subunit, DNA Damage Recognition and Repair Factor

## Grant support

This work was supported by the Research and Technology Innovation Fund (KTIA_NAP_13-2014-0021 and 2017-1.2.1-NKP-2017-00002 to ZS), Breast Cancer Research Foundation (BCRF-20-159 to ZS), Kræftens Bekæmpelse (R281-A16566 to ZS), the Novo Nordisk Foundation Interdisciplinary Synergy Programme Grant (NNF15OC0016584 to ZS), Department of Defense through the Prostate Cancer Research Program (W81XWH-18-2-0056 to ZS), Det Frie Forskningsråd Sundhed og Sygdom (7016-00345B to ZS), the Velux Foundation (00018310 to Zs.S. and JB), the Degregorio Family Foundation (to NSS), AGA Augustyn Award in Digestive Diseases (AGA2021-3101 to NSS), and the NIDDK-National Institute of Health (1K08DK120930 to NSS).

## Disclosure

HS and BMH reveal no conflict of interest. JMC received research funding to his institution from Merus, Roche, and Bristol Myers Squibb. He received research funding from Merck, Astrazeneca, Arcus Biosciences, Apexigen, Esperas Pharma, Bayer, and Tesaro; received advisory board honorarium from Syros Pharmaceuticals and Blueprint Medicines. Z. Szallasi is an inventor on a patent used in the myChoice HRD assay. NS. Sethi is a consultant for Astrin Biosciences.

## Authors contribution

AGP and PS contributed equally. AGP, PS, JB, M.D., Zs.S., Z.S., and NSS made substantial contributions to the conception of the work. PS, CXM, JB, M.D., Zs.S., and AP designed and performed analyses. PS, JB, M.D., Zs.S., Z.S., and NSS wrote and edited the manuscript. P.S., Zs.S., and NSS, substantively revised the manuscript. ZS and NSS supervised the project.

## Data Transparency Statement

All the sequencing data and reagents (e.g. cell lines, inhibitors, antibodies) will be shared either by depositing in public domains or through MTA agreements in compliance with our institutions. There are no restrictions to access the custom code used for the analyses presented in this study. Information is available from the authors on request.

## GRAPHICAL ABSTRACT

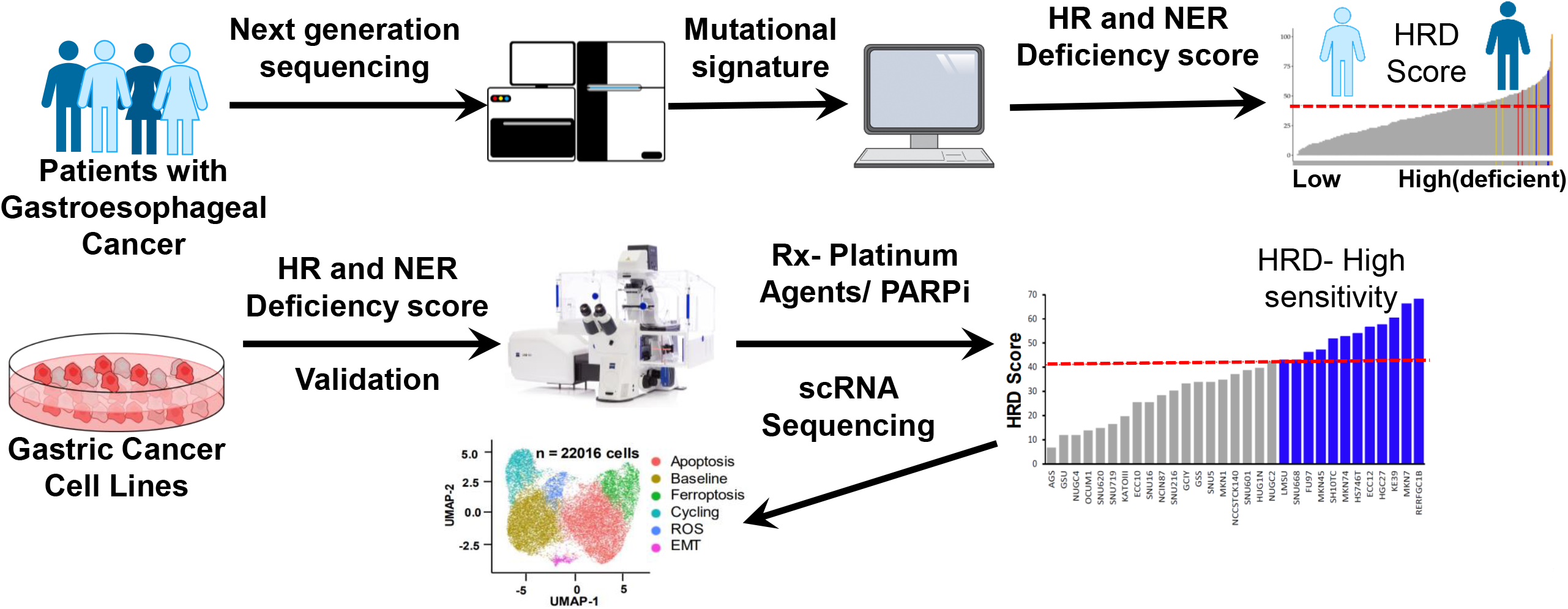

## REFERENCES

1. Silver DP, Richardson AL, Eklund AC, et al. Efficacy of neoadjuvant Cisplatin in triple-negative breast cancer. J Clin Oncol 2010;28:1145–53.

2. Bolton KL, Chenevix-Trench G, Goh C, et al. Association between BRCA1 and BRCA2 mutations and survival in women with invasive epithelial ovarian cancer. JAMA 2012;307:382–90.

3. Cancer Genome Atlas Research N. Comprehensive molecular characterization of gastric adenocarcinoma. Nature 2014;513:202–9.

4. Liu Y, Sethi NS, Hinoue T, et al. Comparative Molecular Analysis of Gastrointestinal Adenocarcinomas. Cancer Cell 2018;33:721–735 e8.

5. Bang YJ, Xu RH, Chin K, et al. Olaparib in combination with paclitaxel in patients with advanced gastric cancer who have progressed following first-line therapy (GOLD): a double-blind, randomised, placebo-controlled, phase 3 trial. Lancet Oncol 2017;18:1637–1651.

6. Bang YJ, Im SA, Lee KW, et al. Randomized, Double-Blind Phase II Trial With Prospective Classification by ATM Protein Level to Evaluate the Efficacy and Tolerability of Olaparib Plus Paclitaxel in Patients With Recurrent or Metastatic Gastric Cancer. J Clin Oncol 2015;33:3858–65.

7. Mirza MR, Coleman RL, Gonzalez-Martin A, et al. The forefront of ovarian cancer therapy: update on PARP inhibitors. Ann Oncol 2020;31:1148–1159.

8. Telli ML, Timms KM, Reid J, et al. Homologous Recombination Deficiency (HRD) Score Predicts Response to Platinum-Containing Neoadjuvant Chemotherapy in Patients with Triple-Negative Breast Cancer. Clin Cancer Res 2016;22:3764–73.

9. Ray-Coquard I, Pautier P, Pignata S, et al. Olaparib plus Bevacizumab as First-Line Maintenance in Ovarian Cancer. N Engl J Med 2019;381:2416–2428.

10. Borcsok J, Sztupinszki Z, Bekele R, et al. Identification of a Synthetic Lethal Relationship between Nucleotide Excision Repair Deficiency and Irofulven Sensitivity in Urothelial Cancer. Clin Cancer Res 2021;27:2011–2022.

11. Goldman MJ, Craft B, Hastie M, et al. Visualizing and interpreting cancer genomics data via the Xena platform. Nat Biotechnol 2020;38:675–678.

12. Favero F, Joshi T, Marquard AM, et al. Sequenza: allele-specific copy number and mutation profiles from tumor sequencing data. Ann Oncol 2015;26:64–70.

13. Sztupinszki Z, Diossy M, Krzystanek M, et al. Migrating the SNP array-based homologous recombination deficiency measures to next generation sequencing data of breast cancer. NPJ Breast Cancer 2018;4:16.

14. Dreze M, Calkins AS, Galicza J, et al. Monitoring repair of UV-induced 6-4-photoproducts with a purified DDB2 protein complex. PLoS One 2014;9:e85896.

15. Rajkumar-Calkins AS, Szalat R, Dreze M, et al. Functional profiling of nucleotide Excision repair in breast cancer. DNA Repair (Amst) 2019;82:102697.

16. Sztupinszki Z, Diossy M, Krzystanek M, et al. Detection of Molecular Signatures of Homologous Recombination Deficiency in Prostate Cancer with or without BRCA1/2 Mutations. Clin Cancer Res 2020;26:2673–2680.

17. Kandoth C, McLellan MD, Vandin F, et al. Mutational landscape and significance across 12 major cancer types. Nature 2013;502:333–339.

18. Sakai W, Swisher EM, Karlan BY, et al. Secondary mutations as a mechanism of cisplatin resistance in BRCA2-mutated cancers. Nature 2008;451:1116–20.

19. Poti A, Gyergyak H, Nemeth E, et al. Correlation of homologous recombination deficiency induced mutational signatures with sensitivity to PARP inhibitors and cytotoxic agents. Genome Biol 2019;20:240.

20. Abkevich V, Timms KM, Hennessy BT, et al. Patterns of genomic loss of heterozygosity predict homologous recombination repair defects in epithelial ovarian cancer. Br J Cancer 2012;107:1776–82.

21. Popova T, Manie E, Rieunier G, et al. Ploidy and large-scale genomic instability consistently identify basal-like breast carcinomas with BRCA1/2 inactivation. Cancer Res 2012;72:5454–62.

22. Birkbak NJ, Wang ZC, Kim JY, et al. Telomeric allelic imbalance indicates defective DNA repair and sensitivity to DNA-damaging agents. Cancer Discov 2012;2:366–375.

23. Boot A, Liu M, Stantial N, et al. Recurrent mutations in topoisomerase IIalpha cause a previously undescribed mutator phenotype in human cancers. Proc Natl Acad Sci U S A 2022;119.

24. Zarrizi R, Higgs MR, Vossgrone K, et al. Germline RBBP8 variants associated with early-onset breast cancer compromise replication fork stability. J Clin Invest 2020;130:4069–4080.

25. Farmer H, McCabe N, Lord CJ, et al. Targeting the DNA repair defect in BRCA mutant cells as a therapeutic strategy. Nature 2005;434:917–21.

26. Robson M, Im SA, Senkus E, et al. Olaparib for Metastatic Breast Cancer in Patients with a Germline BRCA Mutation. N Engl J Med 2017;377:523–533.

27. Gout J, Perkhofer L, Morawe M, et al. Synergistic targeting and resistance to PARP inhibition in DNA damage repair-deficient pancreatic cancer. Gut 2021;70:743–760.

28. Corsello SM, Nagari RT, Spangler RD, et al. Discovering the anti-cancer potential of non-oncology drugs by systematic viability profiling. Nat Cancer 2020;1:235–248.

29. Rose M, Burgess JT, O’Byrne K, et al. PARP Inhibitors: Clinical Relevance, Mechanisms of Action and Tumor Resistance. Front Cell Dev Biol 2020;8:564601.

30. Wang Y, Zheng K, Huang Y, et al. PARP inhibitors in gastric cancer: beacon of hope. J Exp Clin Cancer Res 2021;40:211.

31. Shen Y, Rehman FL, Feng Y, et al. BMN 673, a novel and highly potent PARP1/2 inhibitor for the treatment of human cancers with DNA repair deficiency. Clin Cancer Res 2013;19:5003–15.

32. Oplustil O’Connor L, Rulten SL, Cranston AN, et al. The PARP Inhibitor AZD2461 Provides Insights into the Role of PARP3 Inhibition for Both Synthetic Lethality and Tolerability with Chemotherapy in Preclinical Models. Cancer Res 2016;76:6084–6094.

33. Yang W, Soares J, Greninger P, et al. Genomics of Drug Sensitivity in Cancer (GDSC): a resource for therapeutic biomarker discovery in cancer cells. Nucleic Acids Res 2013;41:D955–61.

34. Marteijn JA, Lans H, Vermeulen W, et al. Understanding nucleotide excision repair and its roles in cancer and ageing. Nat Rev Mol Cell Biol 2014;15:465–81.

35. Li Q, Damish AW, Frazier Z, et al. ERCC2 Helicase Domain Mutations Confer Nucleotide Excision Repair Deficiency and Drive Cisplatin Sensitivity in Muscle-Invasive Bladder Cancer. Clin Cancer Res 2019;25:977–988.

36. Ray A, Milum K, Battu A, et al. NER initiation factors, DDB2 and XPC, regulate UV radiation response by recruiting ATR and ATM kinases to DNA damage sites. DNA Repair (Amst) 2013;12:273–83.

37. Pajuelo-Lozano N, Bargiela-Iparraguirre J, Dominguez G, et al. XPA, XPC, and XPD Modulate Sensitivity in Gastric Cisplatin Resistance Cancer Cells. Front Pharmacol 2018;9:1197.

38. Zhang Y, Cao J, Meng Y, et al. Overexpression of xeroderma pigmentosum group C decreases the chemotherapeutic sensitivity of colorectal carcinoma cells to cisplatin. Oncol Lett 2018;15:6336–6344.

39. Wang C, Nie H, Li Y, et al. The study of the relation of DNA repair pathway genes SNPs and the sensitivity to radiotherapy and chemotherapy of NSCLC. Sci Rep 2016;6:26526.

40. Zhang Y, Yu JJ, Tian Y, et al. eIF3a improve cisplatin sensitivity in ovarian cancer by regulating XPC and p27Kip1 translation. Oncotarget 2015;6:25441–51.

41. Hollander MC, Philburn RT, Patterson AD, et al. Deletion of XPC leads to lung tumors in mice and is associated with early events in human lung carcinogenesis. Proc Natl Acad Sci U S A 2005;102:13200–5.

42. Robu M, Shah RG, Petitclerc N, et al. Role of poly(ADP-ribose) polymerase-1 in the removal of UV-induced DNA lesions by nucleotide excision repair. Proc Natl Acad Sci U S A 2013;110:1658–63.

43. Lord CJ, McDonald S, Swift S, et al. A high-throughput RNA interference screen for DNA repair determinants of PARP inhibitor sensitivity. DNA Repair (Amst) 2008;7:2010–9.

44. Postel-Vinay S, Bajrami I, Friboulet L, et al. A high-throughput screen identifies PARP1/2 inhibitors as a potential therapy for ERCC1-deficient non-small cell lung cancer. Oncogene 2013;32:5377–87.

45. Fleury H, Carmona E, Morin VG, et al. Cumulative defects in DNA repair pathways drive the PARP inhibitor response in high-grade serous epithelial ovarian cancer cell lines. Oncotarget 2017;8:40152–40168.

46. Uemura T, Yerushalmi HF, Tsaprailis G, et al. Identification and characterization of a diamine exporter in colon epithelial cells. J Biol Chem 2008;283:26428–35.

47. Dixon SJ, Lemberg KM, Lamprecht MR, et al. Ferroptosis: an iron-dependent form of nonapoptotic cell death. Cell 2012;149:1060–72.

48. Wu F, Xiong G, Chen Z, et al. SLC3A2 inhibits ferroptosis in laryngeal carcinoma via mTOR pathway. Hereditas 2022;159:6.

49. Mirza MR, Monk BJ, Herrstedt J, et al. Niraparib Maintenance Therapy in Platinum-Sensitive, Recurrent Ovarian Cancer. N Engl J Med 2016;375:2154–2164.

50. Coleman RL, Fleming GF, Brady MF, et al. Veliparib with First-Line Chemotherapy and as Maintenance Therapy in Ovarian Cancer. N Engl J Med 2019;381:2403–2415.

51. Spivak G. Nucleotide excision repair in humans. DNA Repair (Amst) 2015;36:13–18.

52. Scharer OD. Nucleotide excision repair in eukaryotes. Cold Spring Harb Perspect Biol 2013;5:a012609.

