## Supplementary File for "Diagnostic tool to identify and treat DNA repair deficient gastroesophageal adenocarcinomas"

### Diagnostic detection and treatment of DNA repair deficient gastroesophageal adenocarcinomas

#### INVENTORY

Supplemental Material contains 19 Figures and 4 Tables (2 in .xls format)

Supplemental Figure 1-8 is related to Figure 1.
Supplemental Figure 10-13 is related to Figure 2.
Supplemental Figure 14-16 is related to Figure 3.
Supplemental Figure 17 is related to Figure 4.
Supplemental Figure 18-19 is related to Figure 5.

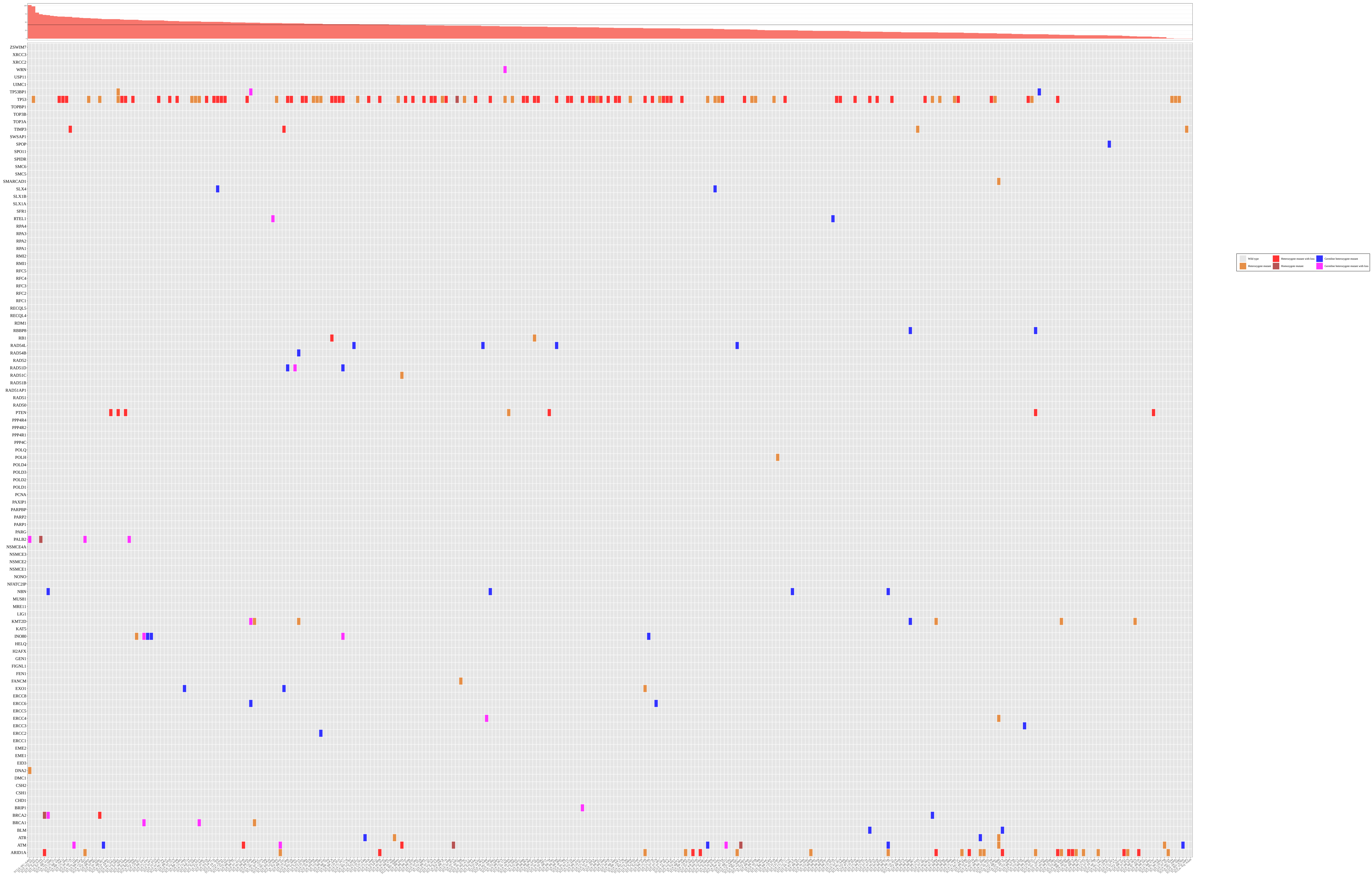

**Supplementary Figure 1| Mutation status of HR genes and DNA damage/DNA-repair response in TCGA-STAD WES.** Cases are ordered by their respective HRD scores.

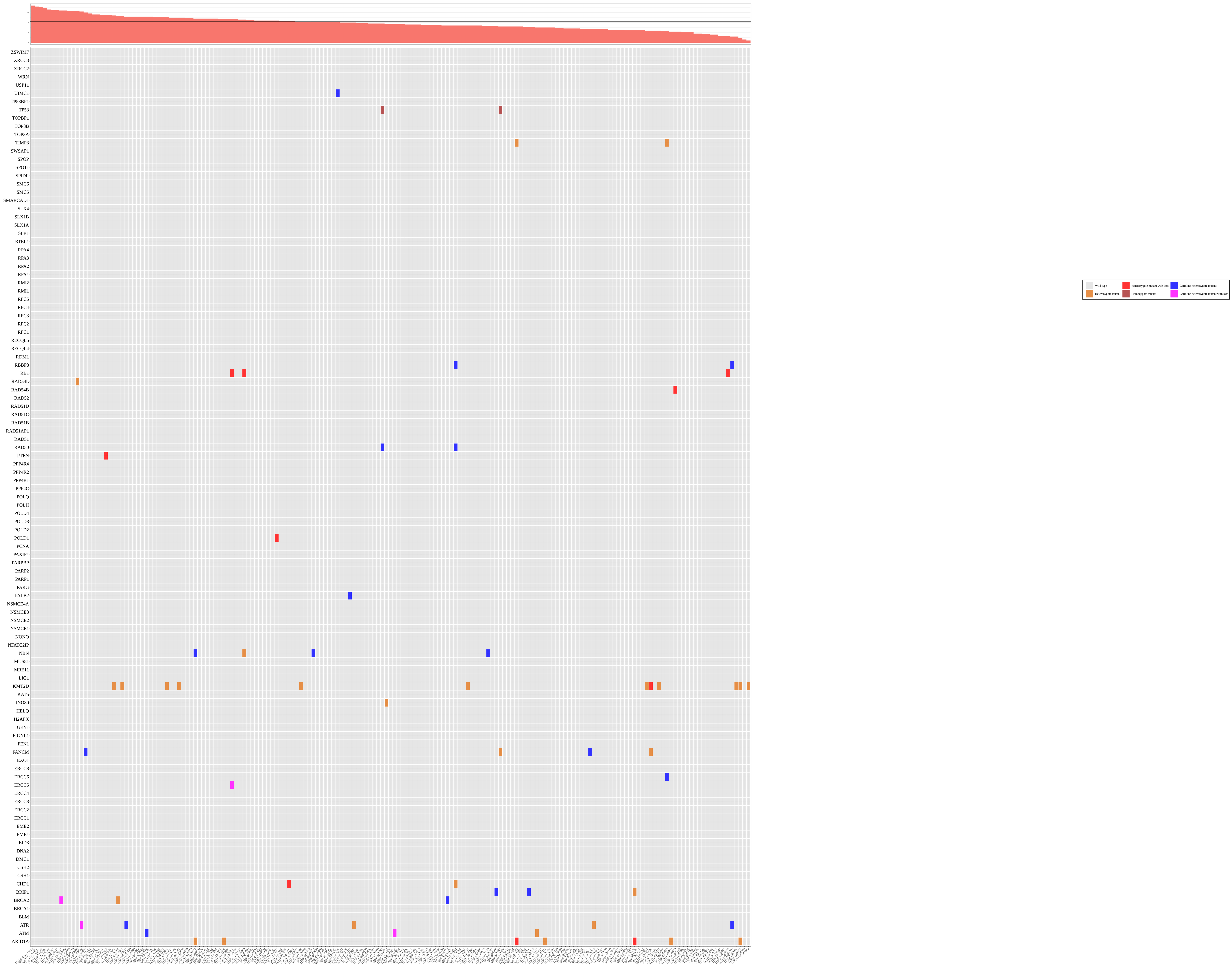

**Supplementary Figure 2| Mutation status of HR genes and DNA damage/DNA-repair response in TCGA-ESCA WES samples.** Cases are ordered by their respective HRD scores.

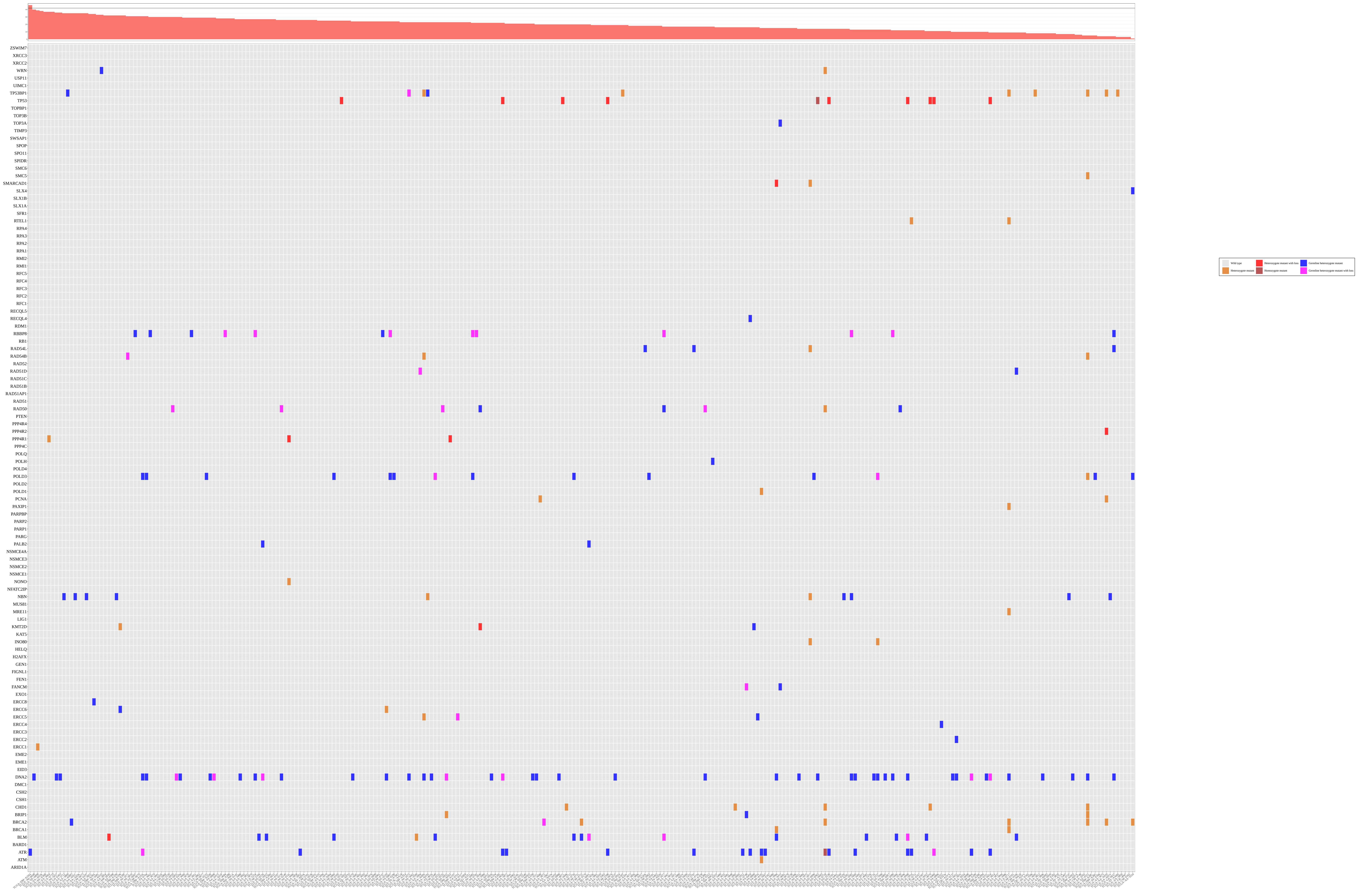

**Supplementary Figure 3| Mutation status of HR genes and DNA damage/DNA-repair response in TCGA-COAD WES samples.** Cases are ordered by their respective HRD scores.

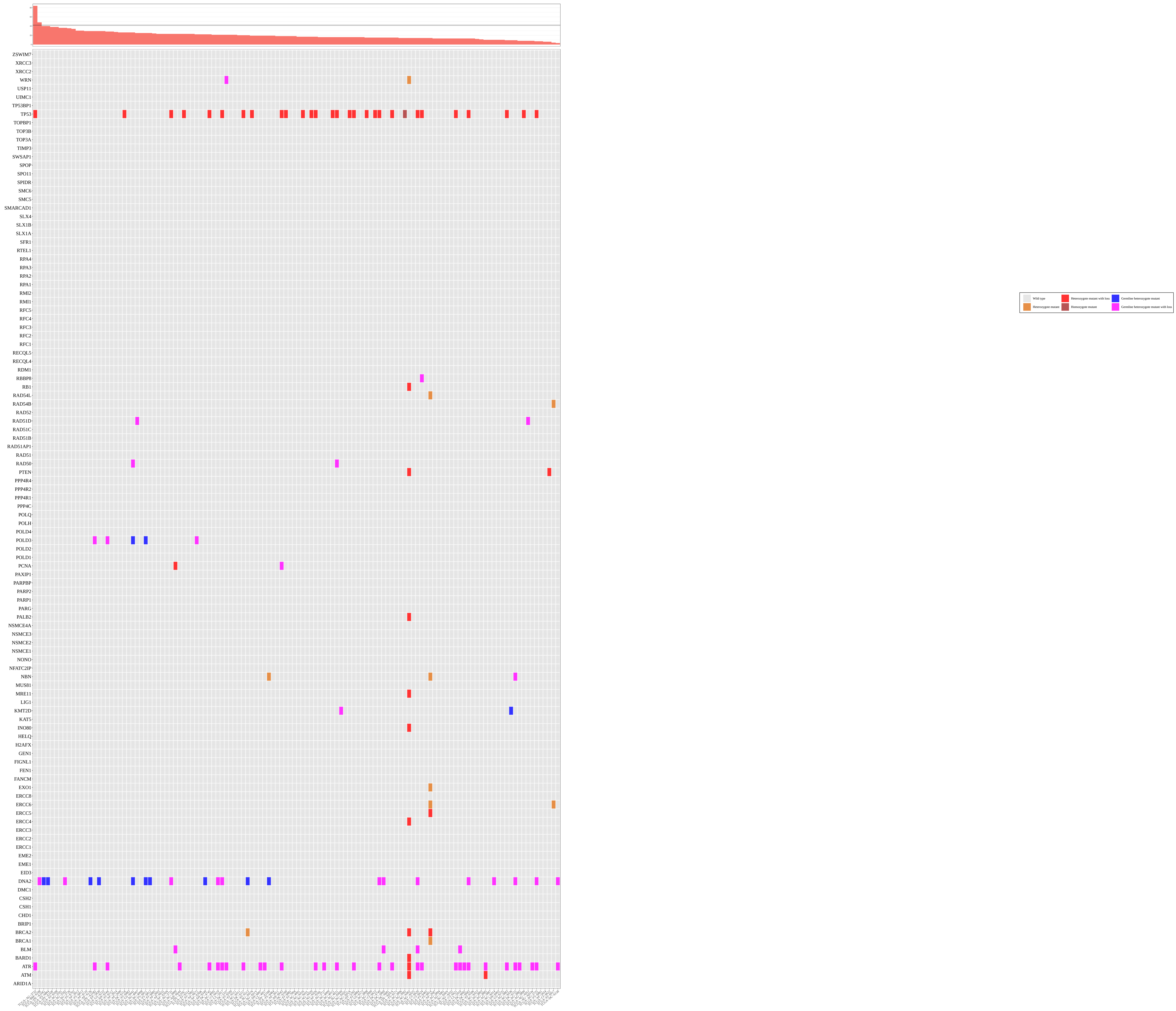


**Supplementary Figure 4| Mutation status of HR genes and DNA damage/DNA-repair response in TCGA-READ WES samples.** Cases are ordered by their respective HRD scores.

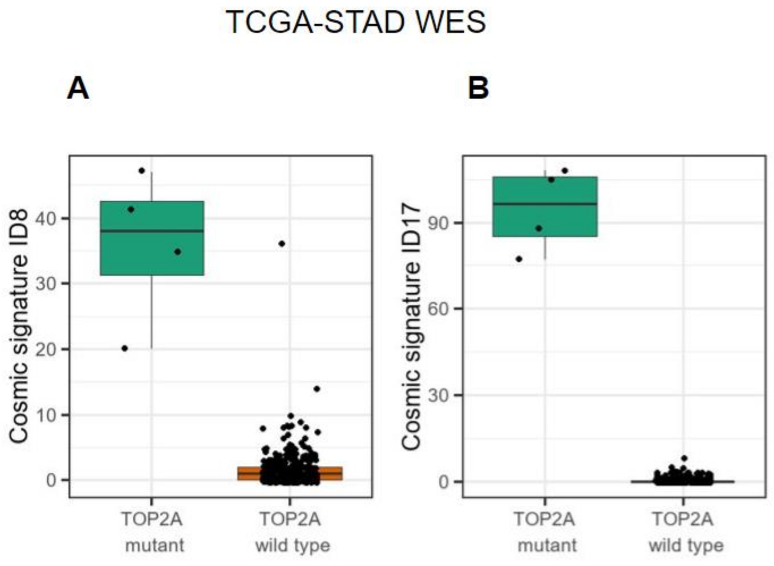


**Supplementary Figure 5| Cosmic signature ID8 and ID17 in the TCGA-STAD WES cohort compared between TOP2A mutants and WT cases.** P-values calculated by Mann-Whitney test.

A. Distribution of Cosmic signature ID8 absolute counts.
B. Distribution of Cosmic signature ID17 absolute counts.

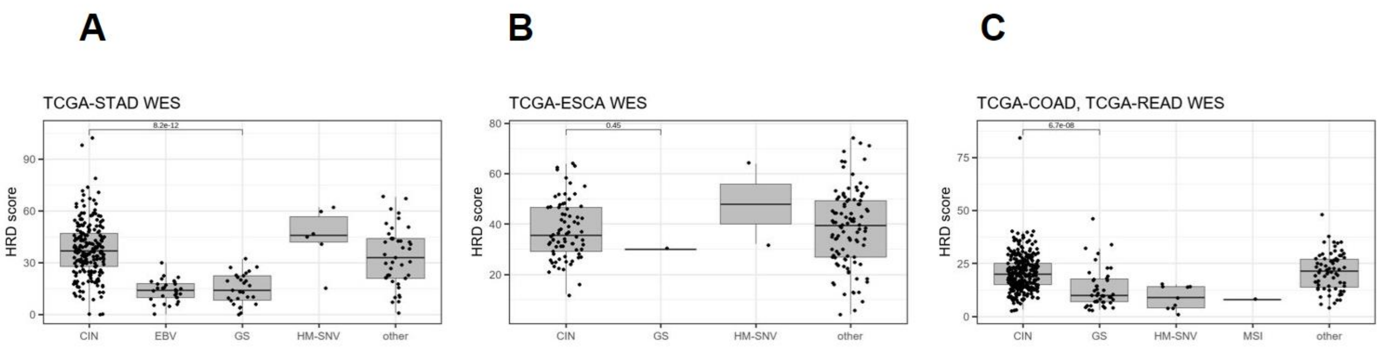


**Supplementary Figure 6| HRD score compared across different molecular subtypes in the TCGA-STAD WES (A), TCGA-ESCA WES (B), TCGA-COAD/READ WES (C) cohorts.** The subtypes are: chromosomal instability (CIN), Epstein–Barr virus (EBV), genomically stable (GS), hypermutated-SNV (HM-SNV). The category other refers to the samples which could not be classified. P-values calculated by Mann Whitney test between the chromosomal instability and genomically stable subtypes.

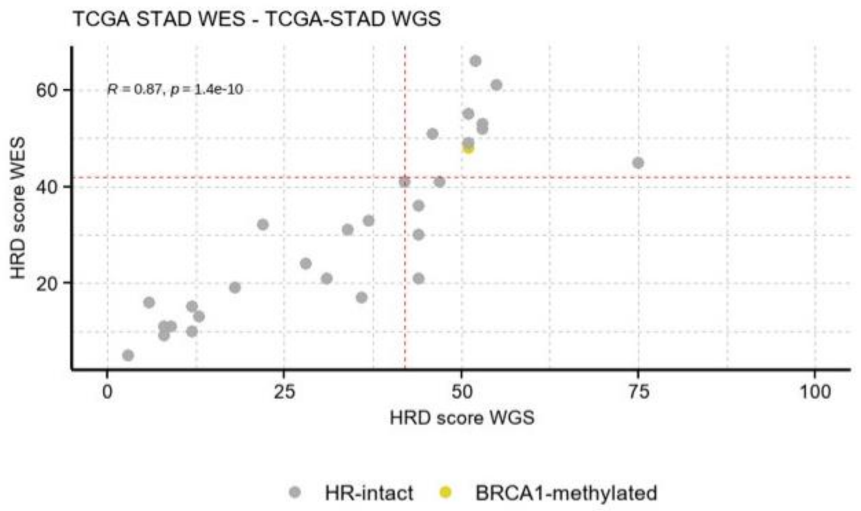


**Supplementary Figure 7| Correlation between the TCGA-STAD WES and TCGA-STAD WGS HRD scores, calculated for the overlapping samples.** Pearson correlation coefficient (0.87), and p-value (1.4e-10) were calculated between the HRD scores of the same TCGA-STAD WES and TCGA-STAD WGS samples and the one BRCA1-methylated case is highlighted.

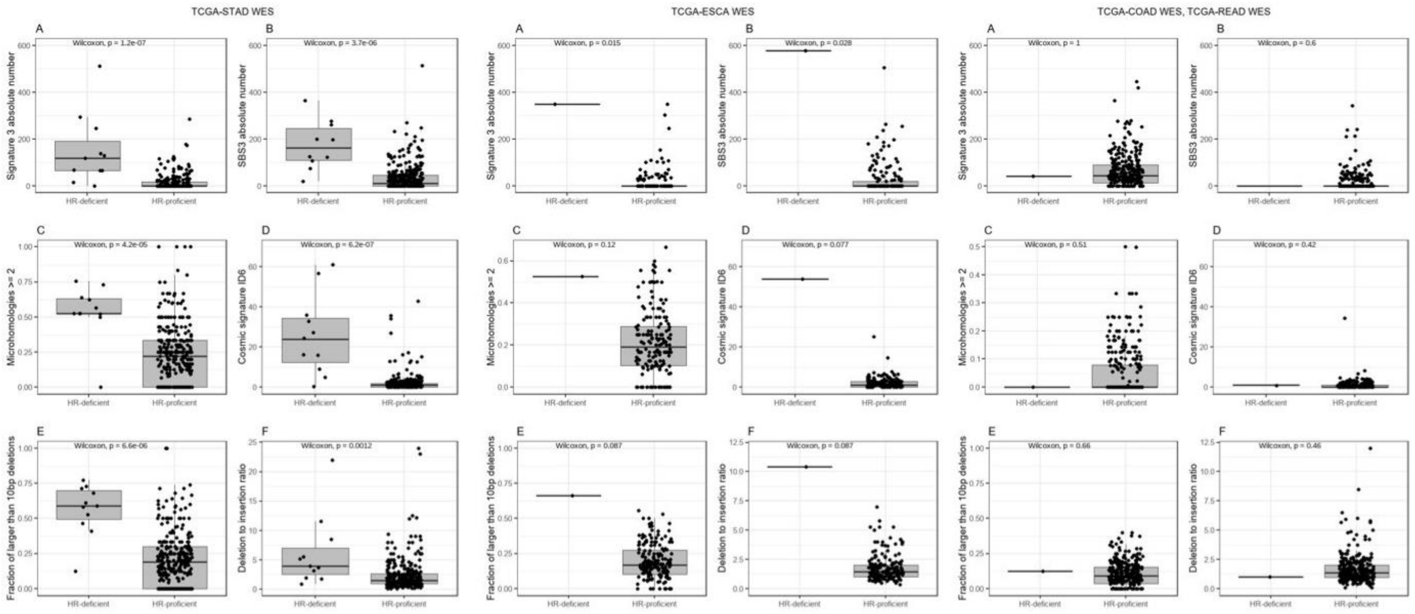


**Supplementary Figure 8| Distribution of signatures associated with BRCA1/2 deficiency in the TCGA STAD WES, TCGA ESCA WES and TCGA COAD/READ WES cohorts.** P-values calculated by Mann Whitney test.

A. Cosmic V2 signature 3 absolute counts.
B. Cosmic V3 SBS3 absolute counts.
C. Fraction of microhomologies at indels, which are equal or longer than 2 base-pairs in length.
D. Cosmic signature ID6 absolute counts.
E. Fraction of larger than 10 base-pair deletions.
F. Deletions to insertion ratio.

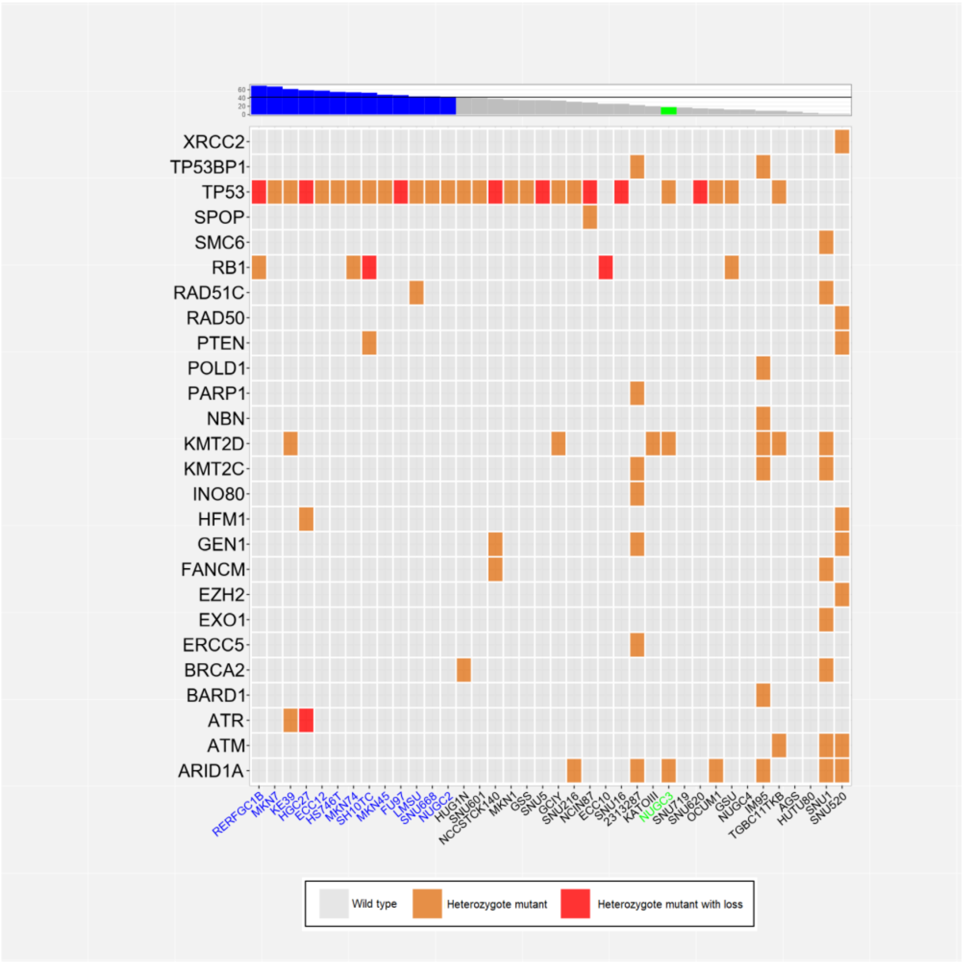


**Supplementary Figure 9| Mutation status of HR genes and DNA damage/DNA-repair response in gastric cancer and non-neoplastic cell lines.** Cell lines are ordered by their respective HRD scores, and they are highlighted in blue, if the HRD score is higher than 42, and NUGC3 is highlighted in green, indicating the functional NER deficiency.

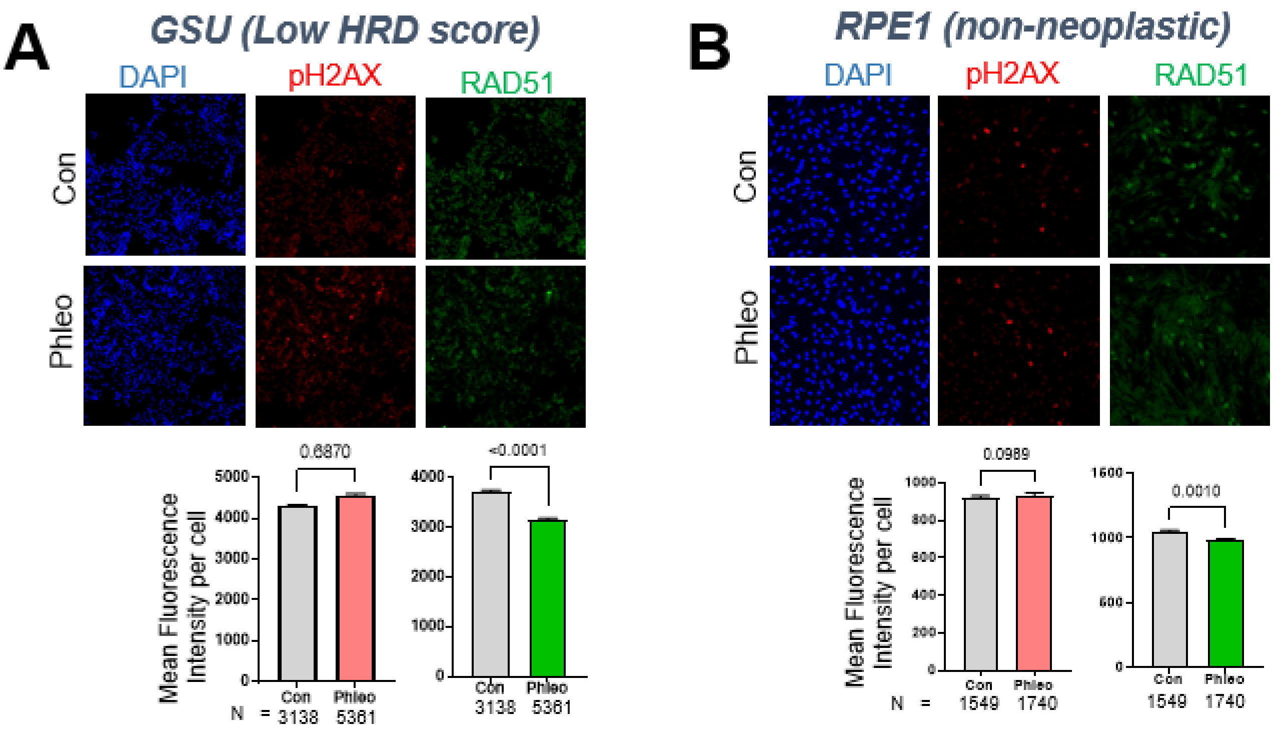


**Supplementary Figure 10| Mutation status of HR genes and DNA damage/DNA-repair response in gastric cancer and non-neoplastic cell lines**

A. Representative immunofluorescence images and quantification from of pH2AX signal (red) indicating DS-DNA breaks and RAD51 signal (green) indicating DS-DNA repair after treatment with DMSO control or radiomemetic (gamma ray) phleomycin; GSU cell line (low HRD score) in grey font; Data expressed as mean fluorescence intensity per cell ± S.D from the number of cells mentioned in the graph for each condition. P-values calculated by Mann Whitney test.

B. Representative immunofluorescence images and quantification from of pH2AX signal (red) indicating DS-DNA breaks and RAD51 signal (green) indicating DS-DNA repair after treatment with DMSO control or radiomemetic (gamma ray) phleomycin; RPE1 cell line (non-neoplastic) in grey font; Data expressed as mean fluorescence intensity per cell ± S.D from the number of cells mentioned in the graph for each condition. P-values calculated by Mann Whitney test.

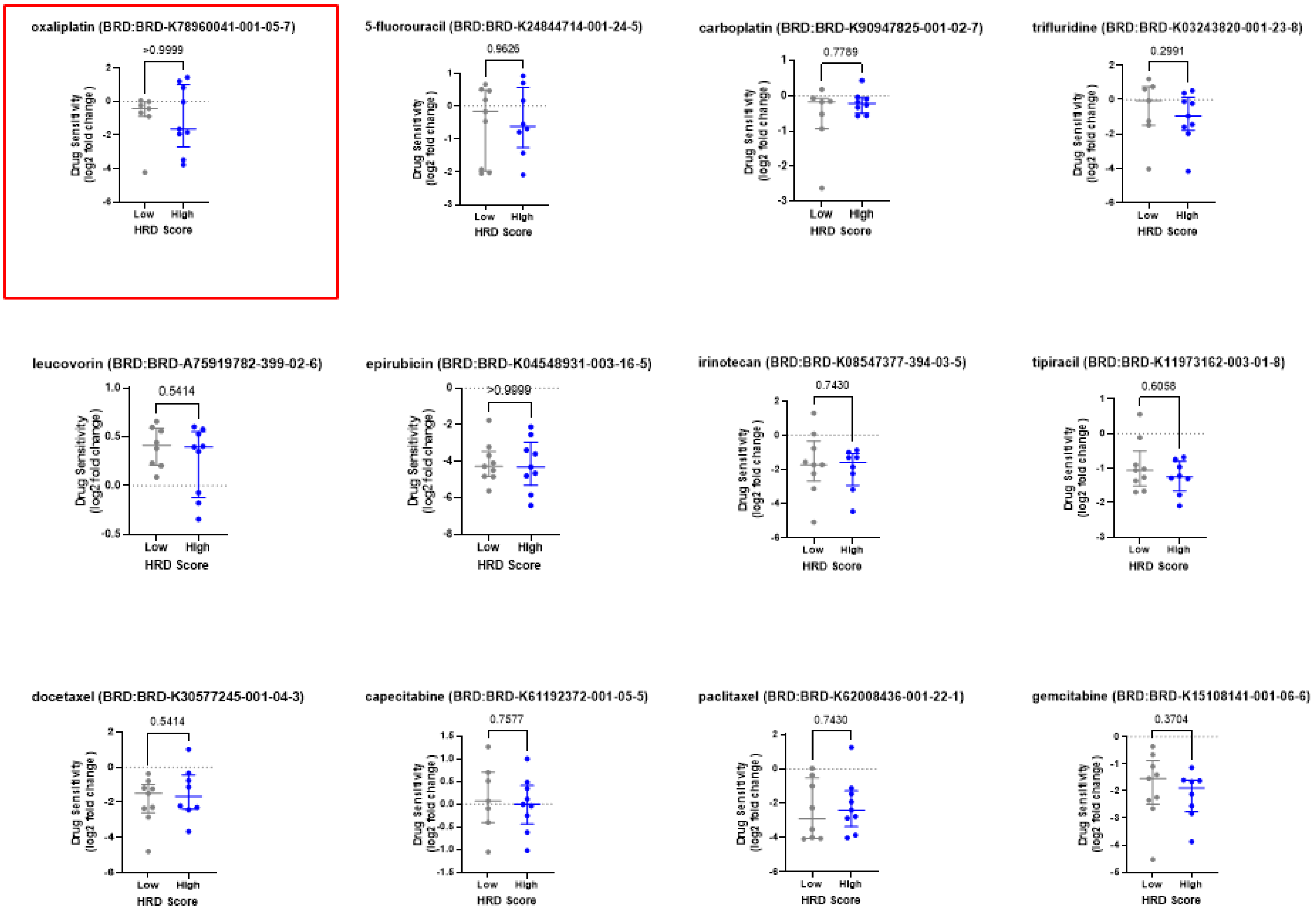


**Supplementary Figure 11| Differential sensitivity of Low (grey) and high (blue) HRD score groups of gastric cancer cell lines for chemotherapeutic agents in BROAD institute PRISM repurposing drug screen dataset**

Commonly used chemotherapeutic agents in GI cancers and other cancers treatment regimen. Difference between the drug sensitivity (log2 fold change) is represented as the median. P-values calculated by Mann Whitney test.

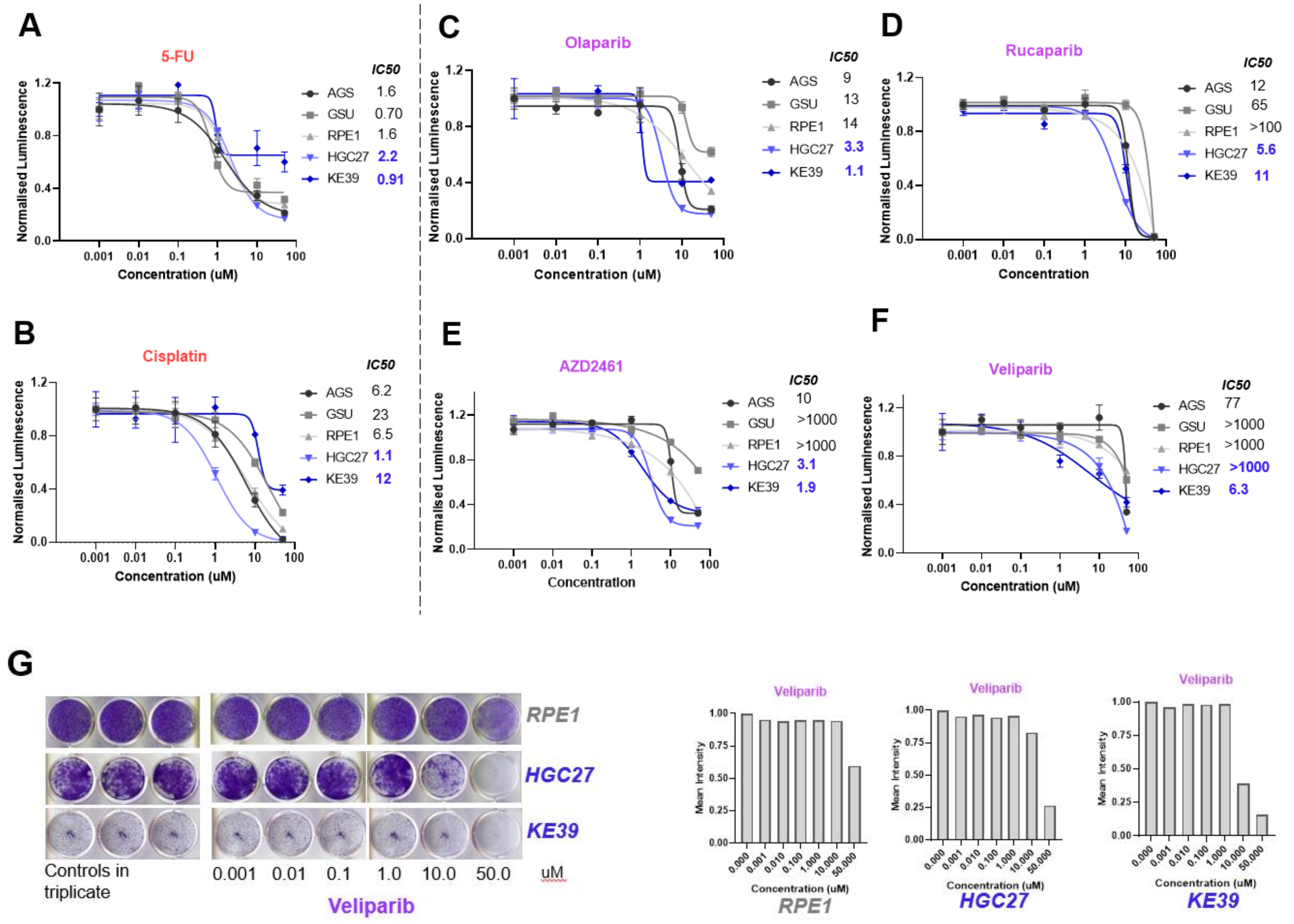


**Supplementary Figure 12| Differential sensitivity of Low (grey) and high (blue) HRD score gastric cell lines for chemotherapeutic agents (red) and PARP inhibitors (magenta)**

A. Dose-response curve of non-neoplastic cell line RPE1, Low HRD (grey) and high HRD (blue) score gastric cancer cell lines to indicated concentrations of 5-FU (red). Best-fit IC50 scores are displayed. Data presented as mean ± s.d. of three culture replicates at each indicated dose.

B. Dose-response curve of non-neoplastic cell line RPE1, Low HRD (grey) and high HRD (blue) score gastric cancer cell lines to indicated concentrations of Cisplatin. Best-fit IC50 scores are displayed. Data presented as mean ± s.d. of three culture replicates at each indicated dose.

C. Dose-response curve of non-neoplastic cell line RPE1, Low HRD (grey) and high HRD (blue) score gastric cancer cell lines to indicated concentrations of Olaparib (magenta). Best-fit IC50 scores are displayed. Data presented as mean ± s.d. of three culture replicates at each indicated dose.

D. Dose-response curve of non-neoplastic cell line RPE1, Low HRD (grey) and high HRD (blue) score gastric cancer cell lines to indicated concentrations of Rucaparib (magenta). Best-fit IC50 scores are displayed. Data presented as mean ± s.d. of three culture replicates at each indicated dose.

E. Dose-response curve of non-neoplastic cell line RPE1, Low HRD (grey) and high HRD (blue) score gastric cancer cell lines to indicated concentrations of AZD2462 (magenta). Best-fit IC50 scores are displayed. Data presented as mean ± s.d. of three culture replicates at each indicated dose.

F. Dose-response curve of non-neoplastic cell line RPE1, Low HRD (grey) and high HRD (blue) score gastric cancer cell lines to indicated concentrations of Veliparib (magenta). Best-fit IC50 scores are displayed. Data presented as mean ± s.d. of three culture replicates at each indicated dose.

G. Colony formation assay (left) and the quantification shown as mean intensity from each well using imageJ (right) of gastric cancer cell with High HRD score (in blue font) and non-neoplastic RPE1 cell line (in grey).

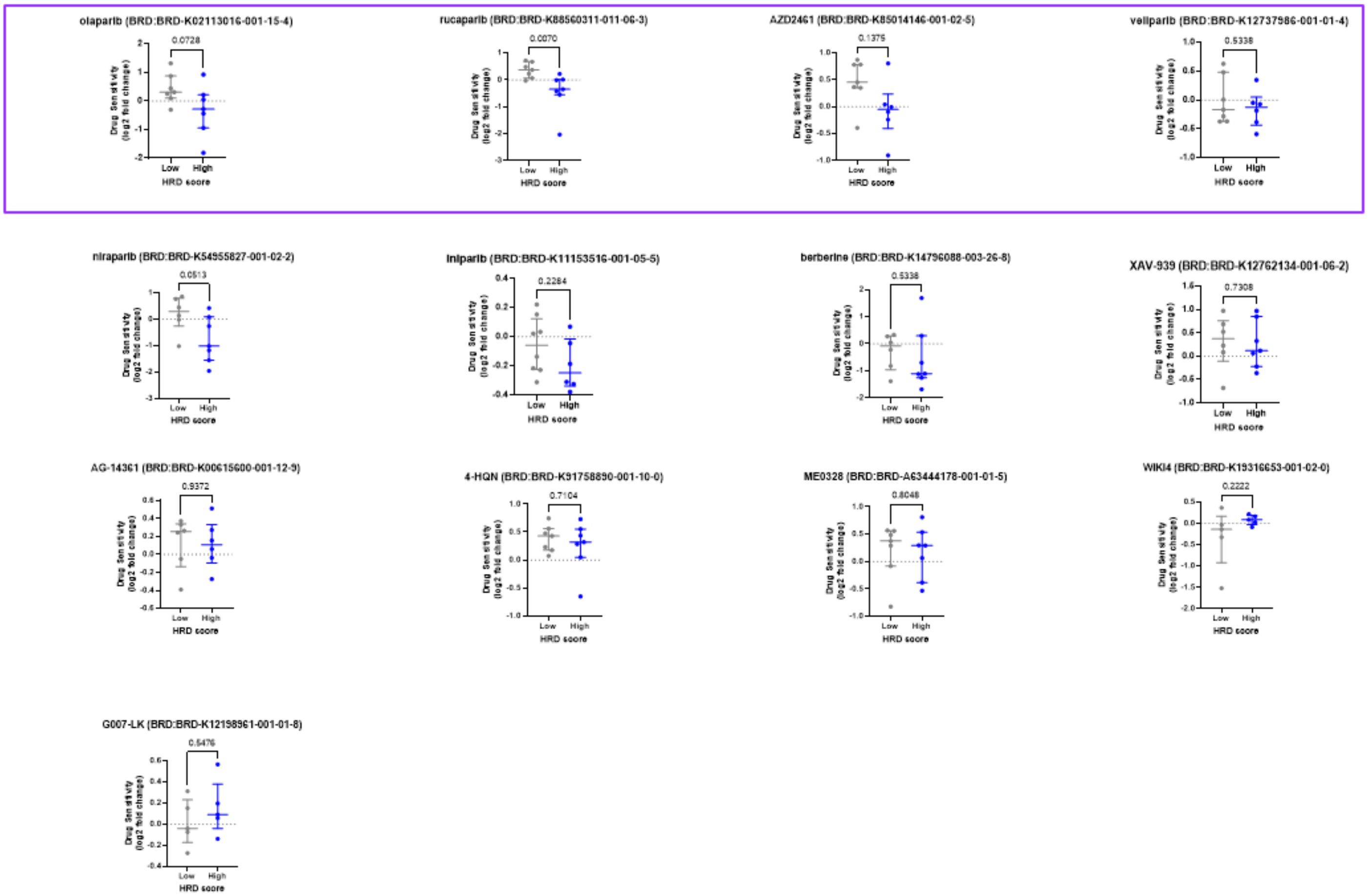


**Supplementary Figure 13| Differential sensitivity of Low (grey) and high (blue) HRD score groups of gastric cancer cell lines for PARP inhibitors in BROAD institute PRISM repurposing drug screen dataset**

Commonly used PARP inhibitors for different cancers. Difference between the drug sensitivity (log2 fold change) is represented as the median. P-values calculated by Mann Whitney test.

PARP inhibitors used in this study are enclosed in magenta box (along with the talazoparib shown in the main figure).

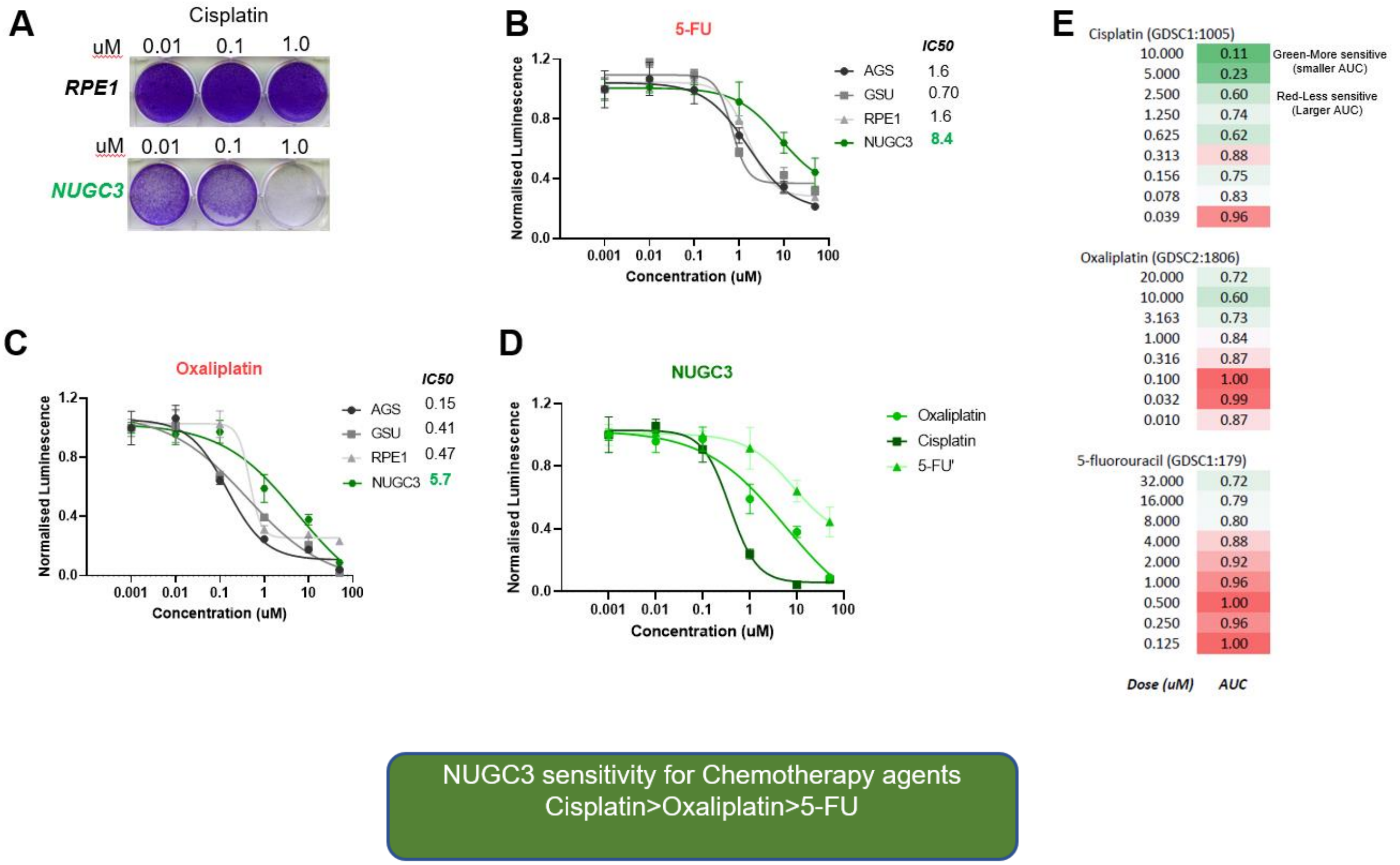


**Supplementary Figure 14| Differential sensitivity of NUGC3 for chemotherapeutic agents**

A. Colony formation assay showing the sensitivity of NUGC3 and RPE1 cell lines for cisplatin at three doses.

B. Dose-response curve of NUGC3 (green) and three control cell lines with intact DNA-repair pathways (grey) to indicated concentrations of chemotherapeutic agent 5-Fu. Best-fit IC50 scores are displayed. Data presented as mean ± s.d. of three culture replicates at each indicated dose.

C. Dose-response curve of NUGC3 (green) and three control cell lines with intact DNA-repair pathways (grey) to indicated concentrations of chemotherapeutic agent Oxaliplatin Best-fit IC50 scores are displayed. Data presented as mean ± s.d. of three culture replicates at each indicated dose.

D. Dose-response curve of NUGC3 cell line to indicated concentrations of chemotherapeutic agents (Oxaliplatin, Cisplatin and 5-Fu). Best-fit IC50 scores are displayed. Data presented as mean ± s.d. of three culture replicates at each indicated dose.

E. NUGC3 cell line checked for Cisplatin, oxaliplatin, and 5-FU sensitivity in Sanger's Genomics of Drug Sensitivity in Cancer Project (GDSC1); Drug response measured as Area Under the Curve (AUC). Dark green color in the heatmap indicates smaller AUC hence higher sensitivity; dark red color indicates larger AUC and relatively lower sensitivity for the chemotherapeutic agent.

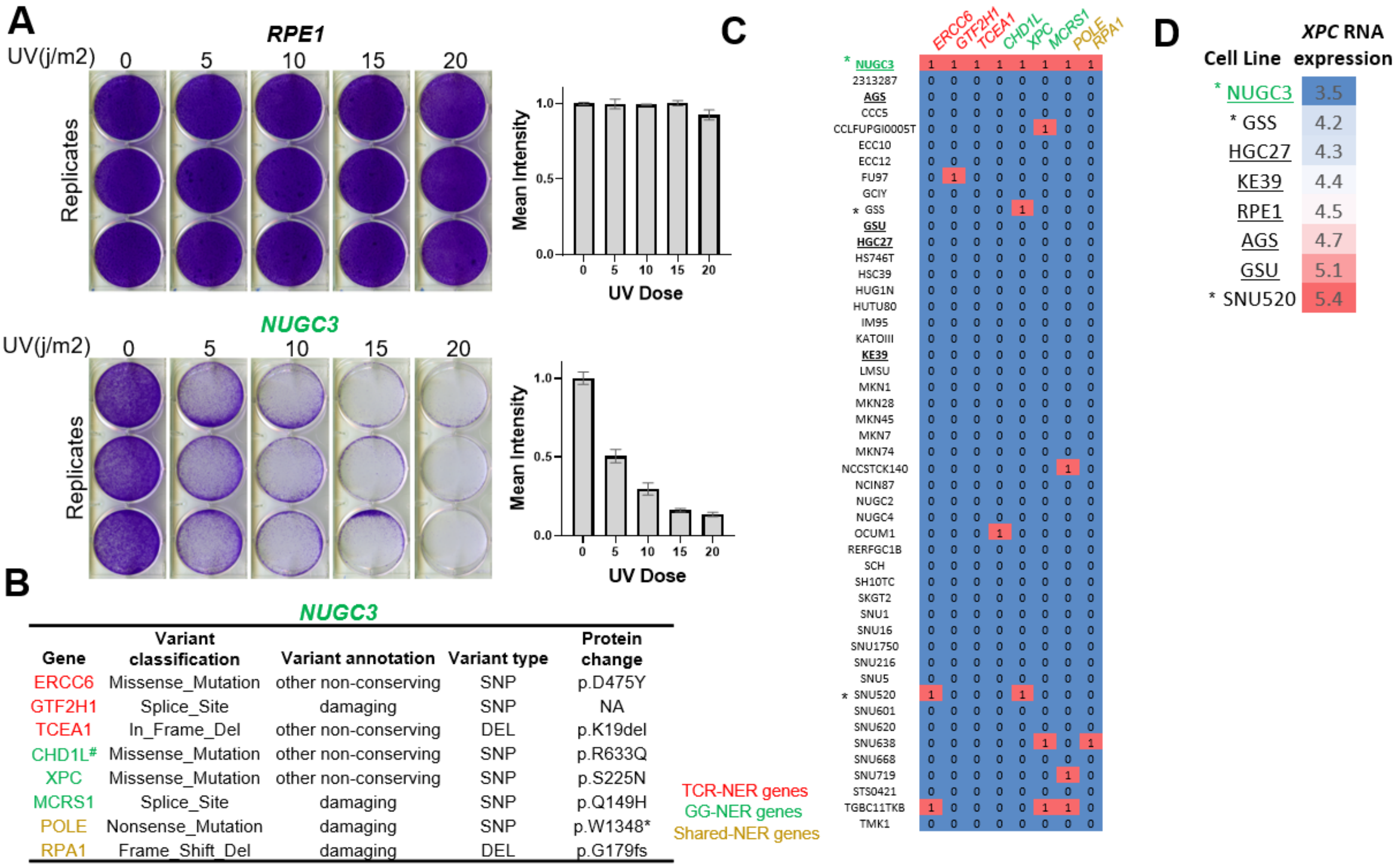


| Gene | Variant classification | Variant annotation | Variant type | Protein change |
|---|---|---|---|---|
| ERCC6 | Missense_Mutation | other non-conserving | SNP | p.D475Y |
| GTF2H1 | Splice_Site | damaging | SNP | NA |
| TCEA1 | In_Frame_Del | other non-conserving | DEL | p.K19del |
| CHD1L# | Missense_Mutation | other non-conserving | SNP | p.R633Q |
| XPC | Missense_Mutation | other non-conserving | SNP | p.S225N |
| MCRS1 | Splice_Site | damaging | SNP | p.Q149H |
| POLE | Nonsense_Mutation | damaging | SNP | p.W1348* |
| RPA1 | Frame_Shift_Del | damaging | DEL | p.G179fs |

**Supplementary Figure 15 | UV-C sensitivity and mutation profile of important NER genes in NUGC3 cell line.**

A. Colony formation assay showing the UV radiation dose-dependent sensitivity (left) and quantification (right) of NUGC3 and RPE1 cell lines. Data presented as mean intensity ± s.d. of three culture replicates at each indicated UV dose.

B. Somatic point mutations and indels in important NER pathway genes in NUGC3 cell line. TCR-NER genes in red, GG-NER genes in green, Shared-NER genes in yellow. Data source: DepMap Internal 22Q1 mutations. #Known pathogenic mutation in breast cancer (COSMIC Genomic Mutation ID COSV63615250).

C. Presence (red/1) or absence (blue/0) of somatic point mutations and indels in important NER pathway genes in all the CCLE gastric cancer cell lines. TCR-NER genes in red, GG-NER genes in green, Shared-NER genes in yellow; NUGC3 in green, Cell lines used in this study are underlined, cell lines with xpc mutation are marked by a star. Data source: DepMap Internal 22Q1 mutations.

D. RNA *XPC* expression TPM values. Dark blue color in the heatmap indicates the lowest expression of xpc; dark red color indicates relatively highest expression of xpc; NUGC3 in green, Cell lines used in this study are underlined, Cell lines with *XPC* mutation are marked by a star. Data source: DepMap Expression 22Q1 Internal.

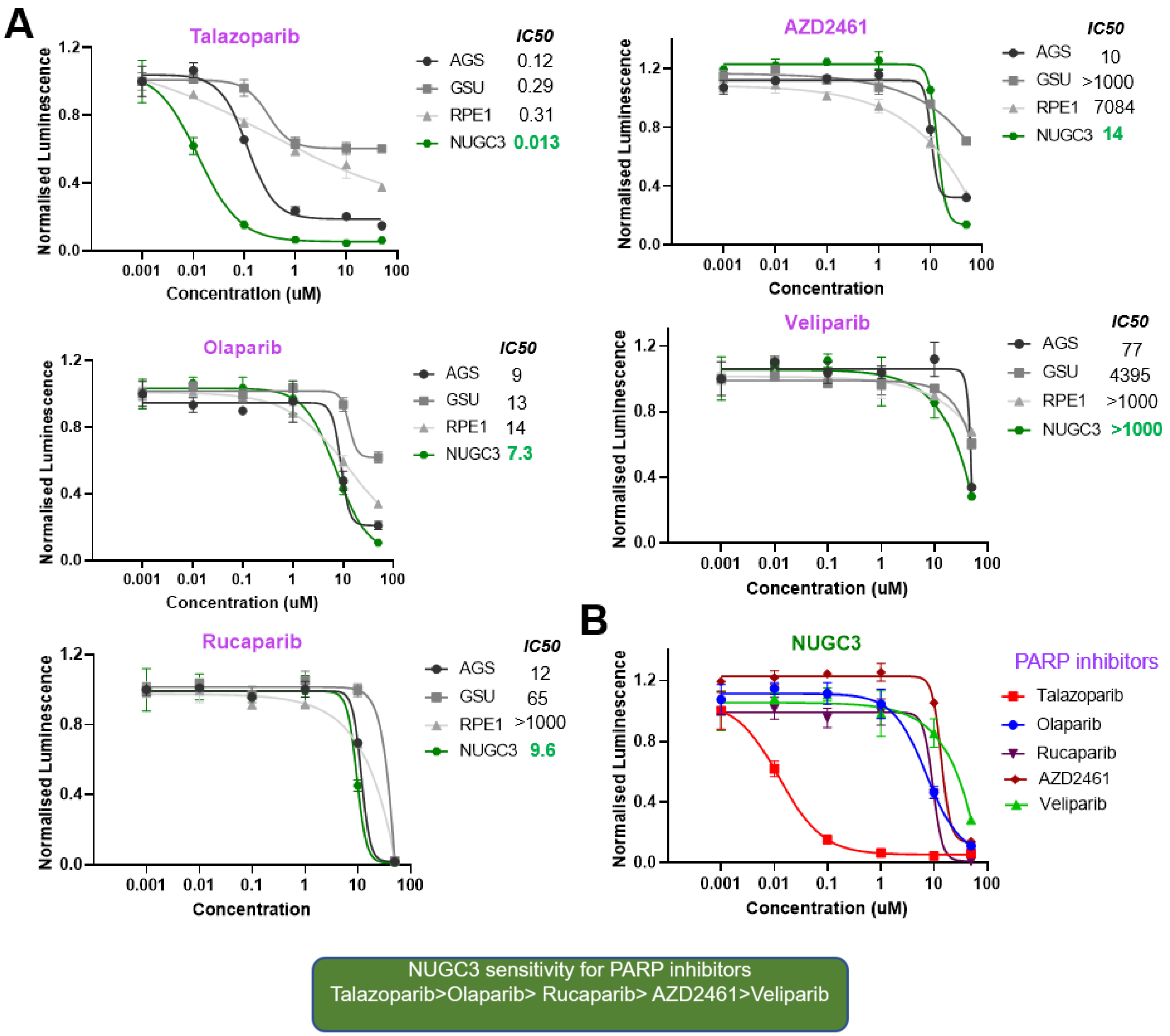


**Supplementary Figure 16| Differential sensitivity of NUGC3 for PARP inhibitors**

A. Dose-response curve of NUGC3 (green) and three control cell lines with intact DNA-repair pathways (grey) to indicated concentrations of PARP inhibitors (Talazoparib, Olaparib, Rucaparib, AZD2461, and Veliparib). Best-fit IC50 scores are displayed. Data presented as mean ± s.d. of three culture replicates at each indicated dose.

B. Dose-response curve of NUGC3 cell line to indicated concentrations of PARP inhibitors (Talazoparib, Olaparib, Rucaparib, AZD2461 and Veliparib). Best-fit IC50 scores are displayed. Data presented as mean ± s.d. of three culture replicates at each indicated dose.

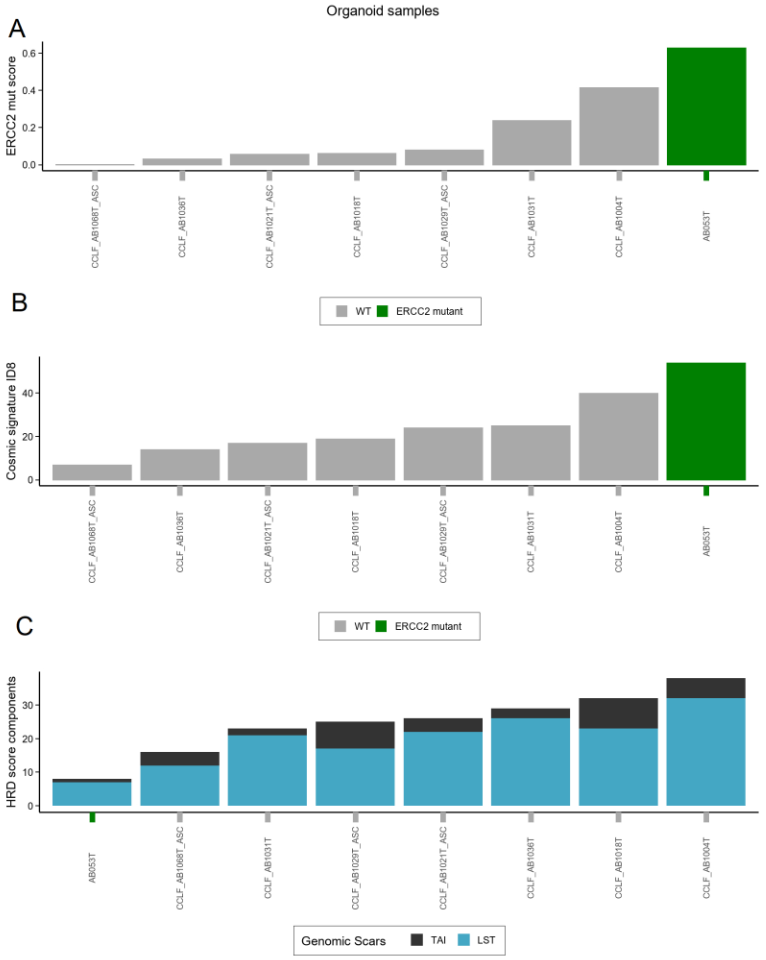


**Supplementary Figure 17| The NER deficiency associated mutational signature score (ERCC2 mut score), Cosmic signature ID8, and two of the HRD score components in the organoid samples**

A. Organoid samples ordered by ERCC2 mut score. The ERCC2 mutant sample is also highlighted here.
B. Organoid samples ordered by Cosmic signature ID8 absolute counts, which is also associated with NER deficiency.
C. Organoid samples ordered by the sum of two of the HRD score components. HRD-LOH was not calculated here, since the normal samples for the organoids were not available.

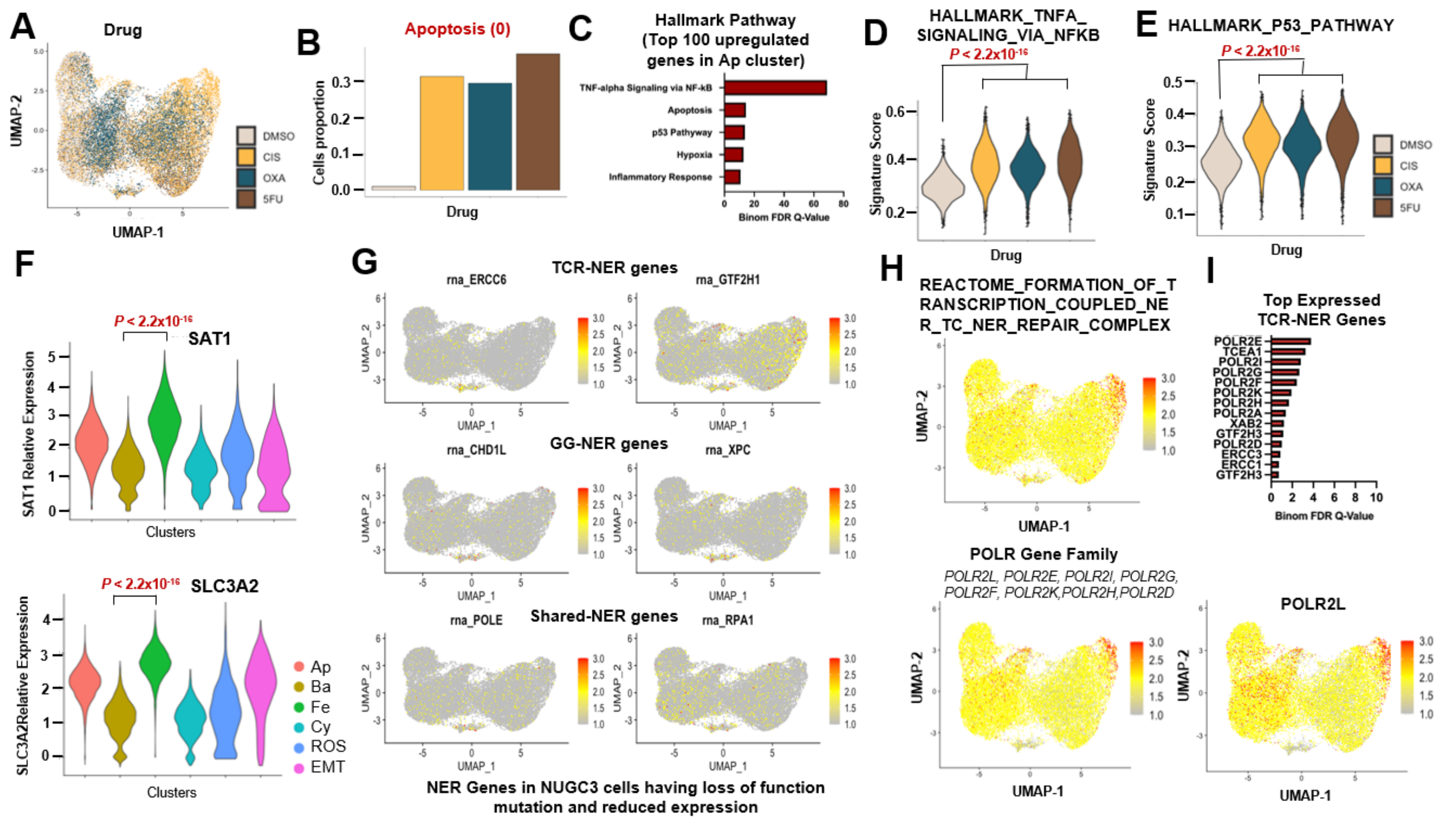


**Supplementary Figure 18 | scRNA-seq analysis of NER deficient gastric cancer cells**

A. UMAP representation of single cell transcriptome profiling of NUGC3 cells treated with DMSO (control) and Cisplatin, Oxaliplatin or 5FU (experimental) colored by and four treatment groups/sample identity.

B. Normalized cell counts in each treatment group for Apoptosis (0) cluster.

C. Hallmark pathways enrichment analysis (EnrichR) of top 100 genes upregulated in Apoptosis (0).

D. Violin plots indicating expression of HALLMARK_TNFA_SIGNALING_VIA_NFKB pathway in the four treatment groups revealed by hallmark pathways enrichment analysis (EnrichR) of top genes upregulated in cluster Ap(0) (also Supp table 1) . P-value calculated by Wilcoxon rank sum test with Bonferroni correction.

E. Violin plots indicating expression of HALLMARK_P53 pathway in the four treatment groups revealed by hallmark pathways enrichment analysis (EnrichR) of top genes upregulated in cluster Ap(0) (also Supp table 1) . P-value calculated by Wilcoxon rank sum test with Bonferroni correction.

F. Violin plots indicating SAT1 (top) and SLC3A2 (bottom) gene expression in different cell clusters. P-value calculated by Wilcoxon rank sum test with Bonferroni correction.

G. UMAP representation of  genes mutated in three subcategory of NER pathways in NUGC3 cells.

H. UMAP representation of TC-NER signature (top), expression of POLR gene family of TC-NER signature (bottom).

I. Top expressed gene members in POLR gene family (top). UMAP representation of POLR2L gene expression (bottom).

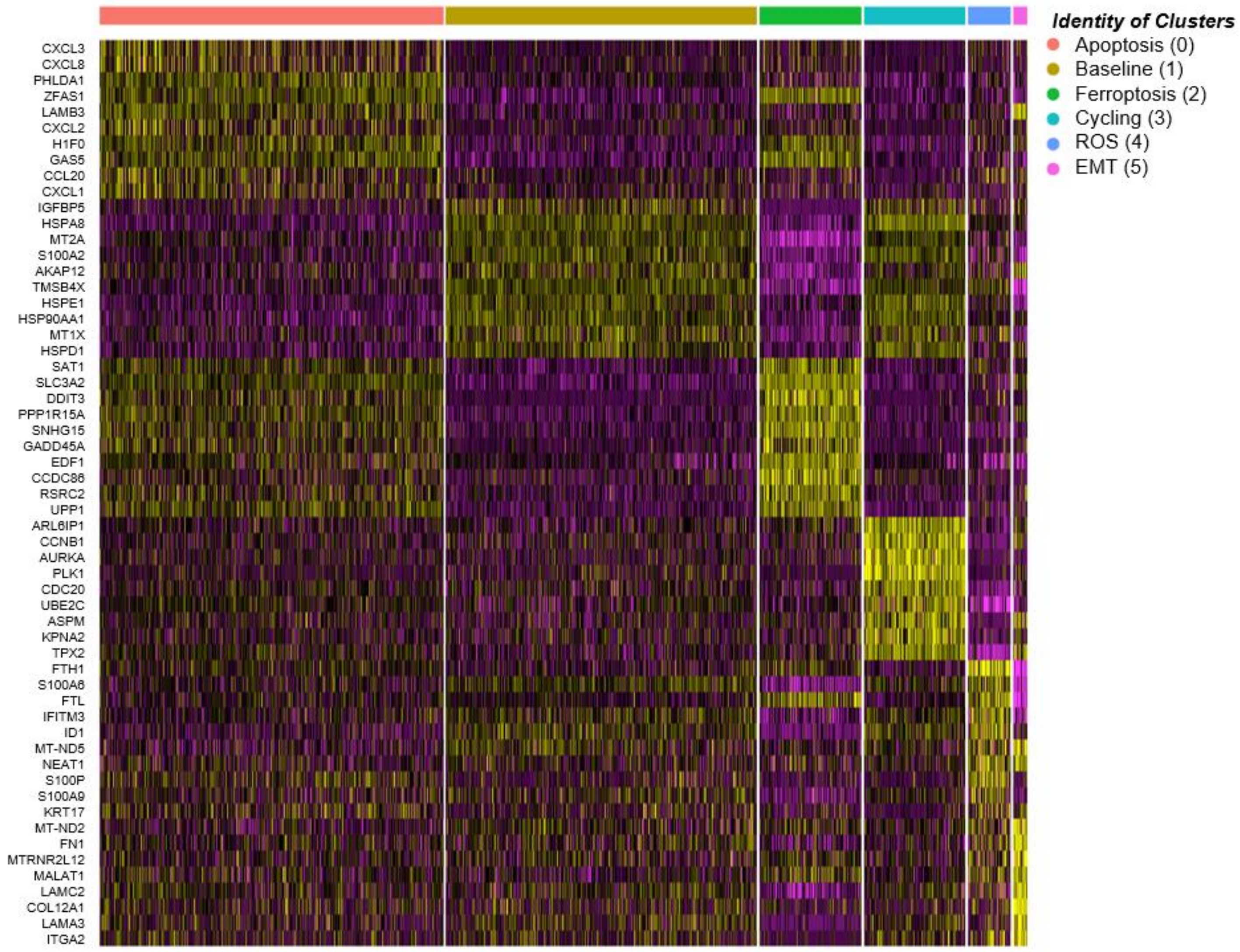


**Supplementary Figure 19 | Heatmap of top 10 differentially expressed genes in each cluster when compared with rest of the clusters.** Dark yellow- highest expression, dark magenta- lowest expression.

**SUPPLEMENTARY TABLES**

| Cohort | Sequencing type | Number of samples |
|---|---|---|
| **TCGA-STAD** | WGS | 32 |
| **ICGC-GACA** | WGS | 36 |
| **ICGC-ESAD** | WGS | 97 |
| **TCGA-COAD** | WGS | 46 |
| **TCGA-READ** | WGS | 16 |
| **TCGA-READ** | WES | 172 |
| **TCGA-COAD** | WES | 461 |
| **TCGA-STAD** | WES | 441 |
| **TCGA-ESCA** | WES | 185 |

**Supplementary Table 1. Number of samples analyzed from each cohort. The actual number of samples presented in the results may differ due to the applied filters.**

| Patient ID | Gene | Chr | POS | REF | ALT | Origin | Function | Genotype | Cohort |
|---|---|---|---|---|---|---|---|---|---|
| **TCGA-BR-4191** | BRCA2 | 13 | 32930687 | C | T | Germline | Stop gain | Heterozygote mutant with LOH | TCGA-STAD |
| **TCGA-BR-4267** | BRCA1 | 17 | 41209080 | - | G | Germline | Frameshift insertion | Heterozygote mutant with LOH | TCGA-STAD |
| **TCGA-D7-A4YT** | BRCA2 | 13 | 32913285 | TCAA | - | Germline | Frameshift deletion | Compound heterozygote mutant | TCGA-STAD |
| **TCGA-D7-A4YT** | BRCA2 | 13 | 32339699 | - | A | Somatic | Frameshift insertion | Compound heterozygote mutant | TCGA-STAD |
| **TCGA-IN-A6RL** | BRCA2 | 13 | 32363286 | C | A | Somatic | Stop gain | Heterozygote mutant with LOH | TCGA-STAD |
| **TCGA-VQ-A91S** | BRCA1 | 17 | 41258474 | T | C | germline | SNP | Heterozygote mutant with LOH | TCGA-STAD |
| **TCGA-L5-A8NS** | BRCA2 | 13 | 32914767 | TT | - | Germline | Frameshift deletion | Heterozygote mutant with LOH | TCGA-ESCA |
| **TCGA-AA-3875** | BRCA2 | 13 | 32954181 | G | A | Germline | SNP | Heterozygote mutant with LOH | TCGA-COAD |
| **TCGA-CG-4437** | PALB2 | 16 | 23649207 | ACAA | - | Germline | Frameshift deletion | Heterozygote mutant with LOH | TCGA-STAD |
| **TCGA-EQ-A4SO** | PALB2 | 16 | 23632683 | C | T | Germline | Stop gain | Heterozygote mutant with LOH | TCGA-STAD |
| **TCGA-HU-A4H5** | PALB2 | 16 | 23640545 | G | A | Germline | Stop gain | Heterozygote mutant with LOH | TCGA-STAD |
| **TCGA-VQ-AA64** | PALB2 | 16 | 23647109 | - | A | Germline | Frameshift insertion | Compound heterozygote mutant | TCGA-STAD |
| **TCGA-VQ-AA64** | PALB2 | 16 | 23629212 | C | A | Somatic | Stop gain | Compound heterozygote mutant | TCGA-STAD |

**Supplementary Table 2. The identified BRCA1/2 and PALB2 deficient cases for the TCGA-STAD WES, TCGA-ESCA WES, TCGA COAD WES and TCGA READ WES cohorts.**
